# Single-Stranded nucleic acid binding enhances the *in vitro* catalytic activity of Chikungunya virus nsP2 protease

**DOI:** 10.1101/2025.05.13.653680

**Authors:** Mohammadamin Mastalipour, Mônika Aparecida Coronado, Ruth Anasthasia Siahaan, Alissa Drees, Christian Ahlers, Markus Fischer, Dieter Willbold, Raphael Josef Eberle

## Abstract

Chikungunya virus (CHIKV) is an emerging arbovirus whose replication relies on the multifunctional nonstructural protein 2 (nsP2), particularly its viral protease (nsP2^pro^), which is essential for polyprotein processing. In this study, we investigated how interactions with nucleic acids influence nsP2^pro^ activity. Using high-throughput sequencing–fluorescent ligand interaction profiling, we identified specific single-stranded DNA aptamers that enhanced nsP2^pro^ activity. Additionally, both random single-stranded DNA and single-stranded RNA were found to stimulate protease activity, whereas double-stranded DNA showed no such effect. Circular dichroism spectroscopy and secondary structure predictions confirmed that the identified aptamers adopt stable folded conformations. Similarly, structured RNA sequences were also capable of promoting protease activity. The observed stimulatory effect depended on the nucleic acid strand type, length, and buffer conditions, suggesting the involvement of electrostatic interactions. Molecular docking analyses further let us assume that these nucleic acids interact specifically with the nsP2^pro^ methyltransferase domain. Our findings provide novel insights into the regulation of nsP2^pro^, enhancing our understanding of CHIKV replication mechanisms, and may guide future antiviral development strategies.

## 1. Introduction

Changes in climate and environmental conditions, including rising temperatures, create favorable environments for the proliferation and spread of vectors such as mosquitoes (Pavia et al., 2025). This, in turn, facilitates the transmission of arthropod-borne viruses to areas where they were previously not endemic, including Chikungunya virus (CHIKV), Dengue virus (DENV), and other tropical infections (Chitre et al., 2024). Chikungunya, a member of the *Alphavirus* genus within the *Togaviridae* family (Alissa, Alsuwat, & Alzahrani, 2024; C. K. Martin et al., 2025), has caused outbreaks across Asia, and South and Central Africa, infected over 3 million people worldwide (Zhang et al., 2025). Approximately 39% of the world’s population lives in endemic regions and remains at risk of infection. (Maneerattanasak et al., 2024). CHIKV can cause a variety of clinical symptoms, including high fever, joint pain, skin rash, and arthralgia, as well as complications such as cardiac and neurological issues (de Souza et al., 2024; Sagar et al., 2024). The infection can progress to a chronic phase characterized by persistent pain and joint inflammation (Silveira-Freitas et al., 2024). Patients in this phase may also experience depression, memory loss, and sleep disturbances (Silveira-Freitas et al., 2024; van Aalst, Nelen, Goorhuis, Stijnis, & Grobusch, 2017), which contribute to disability and significantly reduce quality of life (Resck et al., 2024).

CHIKV is a positive-sense, single-stranded RNA virus whose genome encodes five structural proteins, including envelope (E) and capsid proteins, along with four non-structural proteins (nsP1-nsP4) (Sreekumar et al., 2010; Noranate et al., 2014). Among these, nsP2 is a multifunctional protein essential for viral infection. It contains an N-terminal RNA helicase with both nucleotide triphosphatase (NTPase) and RNA triphosphatase activities, and a C-terminal papain-like cysteine protease domain (nsp2^pro^) responsible for polyprotein processing, followed by a Ftsj methyltransferase (MTase)-like domain (Aher, & Lole, 2011; Das, Merits, & Lulla, 2014). Law et al. 2019 provided a structural analysis of the CHIKV nsP2 helicase domain bound to RNA (Law et al., 2019)

Besides its proteolytic activity, nsP2^pro^ interacts with host proteins, further facilitating viral replication. Notably, studies have demonstrated that nsP2^pro^ migrates to the cell nucleus, although the precise biological significance of this localization remains unclear (Saisawang et al., 2017). The interferon response is a critical first-line defense mechanism of the innate immune system against viral infections, but it can be suppressed by viral factors (Göertz et al., 2018; tenOever, 2016). Specifically, the methyltransferase domain of CHIKV nsP2^pro^ is known to downregulate the interferon response by inhibiting the JAK/STAT signaling pathway, which is essential for interferon activation. Furthermore, this domain disrupts MHC-I antigen presentation, enabling the virus to evade detection by CD8^+^ T cells (Ware et al., 2024). Given that many virus proteases are activated upon interaction with cofactor proteins, these proteases interact with virus and host nucleic acids (DNA or RNA) in several ways, primarily to process viral polyproteins and regulate host cell functions. These interactions are crucial for viral replication, pathogenesis, and immune evasion. Table 1 summarizes reported virus protease-nucleic acid interactions.

**Table 1.** Summary of virus proteases interacting with nucleic acids.

| **Virus** | **Virus protease** | **Nucleic acid** | **Binding/Activation/**<br>**Inhibition** | **Reference** |
| --- | --- | --- | --- | --- |
| **Coxsackievirus** | 3C<sup>pro</sup> | Virus RNA | Binding | (Dias-Solange, Le, Gottipati, & Choi, 2025) |
| **Foamy virus** | Foamy virus protease | Virus RNA | Activation | (Hartl et al., 2011) |
| **Hepatitis A virus** | 3C<sup>pro</sup> | Synthetic DNA | Inhibition | (Blaum et al., 2011) |
| **Hepatitis C virus** | NS3 protease | Virus RNA | Binding | (Beran, Serebrov, & Pyle, 2007) |
| **Hepatitis C virus** | NS3 protease | RNA aptamer | Inhibition | (Urvil et al., 1997) |
| **Human immunodeficiency virus type 1** | HIV-1 PR | Virus RNA | Activation | (Potempa et al., 2015) |
| **Human immunodeficiency virus type 1** | HIV-1 PR | RNA aptamer | Inhibition | (Duclair, Gautam, Ellington, & Prasad, 2015) |
| **Poliovirus** | 3C<sup>pro</sup> | Virus RNA | Activation | (Campagnola & Peersen, 2023) |
| **Seneca Valley virus** | 3C<sup>pro</sup> | Host DNA | Activation | (L. Wu et al., 2025) |

In this study, we explore how nucleic acids influence the catalytic activity of nsP2^pro^ to better understand the regulation of its enzymatic function. Through Throughput Sequencing–Fluorescent Ligand Interaction Profiling (HiTS-FLIP), we identified specific DNA aptamers that bind to nsP2^pro^ and evaluated their effects on its proteolytic activity. Additionally, we demonstrated that random single-stranded RNA (ssRNA) and single-stranded DNA (ssDNA) also modulate nsP2^pro^ function. Finally, *in silico* analyses and molecular docking were conducted to predict the potential binding sites of these nucleic acids on nsP2^pro^.

## 2. Materials and Methods

### 2.1 Expression and purification of nsP2^pro^

The protein was expressed and purified as previously described (Eberle et al., 2021; Mastalipour et al., 2025).

### 2.2 High-Throughput Sequencing–Fluorescent Ligand Interaction Profiling (HiTS-FLIP)

The aptamer selection was performed by HiTS-FLIP as described previously (Drees et al., 2024). In specific, we sequenced a random library (5′-TCGCACATTCCGCTTCTACC-N_50_- CGTAAGTCCGTGTGTGCGAA-3′, acquired PAGE-purified and hand-mixed from Integrated DNA Technologies Inc., Coralville, IA, USA) using a v2 Nano 300 cycle MiSeq kit (Illumina Inc., San Diego, CA, USA) after spiking in 12% phiX (Illumina Inc., San Diego, CA, USA) and 1.25% fiducial mark oligo (acquired from Integrated DNA Technologies Inc., Coralville, IA, USA). Of the 1,682,200 clusters, 84.5% passed filters. For the aptamer selection via HiTS-FLIP, CHIKV nsP2^pro^ was labelled using AF647-NHS-ester (Lumiprobe GmbH, Hannover, Germany), which primarily binds to lysine side chains and the α-amino group at the N-terminus of the protein, with a labelling efficiency of approximately 1 dye molecule per protein. Subsequent to sequencing, the labeled protein was introduced to the flow cell of a modified MiSeq sequencer (Illumina Inc.) at concentrations of 29.8 pM, 149 pM, 745 pM, 3.73 nM, 18.7 nM, 93.2 nM, and 466 nM in 1x PBS + 0.1% Tween20 (pH 7.4, Sigma-Aldrich Corp., St. Louis, MO, USA) via an external valve (C25Z-31812EUHB, Valco Instruments Co. Inc., Houston, TX, USA; VICI). On the flow cell, each protein concentration was incubated for 30 min at 37 °C. For each DNA cluster, fluorescence signals were measured and normalized to reflect the amount of AF647-NHS-ester labelled CHIKV nsP2^pro^ bound at each concentration. Binding curves for each cluster were generated by plotting fluorescence intensity against protein concentration, and K_D_ values were determined using a one-site binding model with Hill-fit analysis in MATLAB R2022b (The MathWorks Inc., Natick, MA, USA). The resulting apparent K_D_ values represent the equilibrium binding affinities under the specific conditions of the flow cell assay (Drees et al., 2024). Out of the 910,150 distinct aptamer library sequences displayed on the flow cell, after filtering the data as described previously (Drees et al., 2024) the ten aptamer candidates with the highest affinity were chosen for further analysis.

### 2.3 Nucleic acids

All DNA aptamers and RNA oligonucleotides were purchased in lyophilized form from Integrated DNA Technologies, Inc. (Coralville, Iowa, USA). DNA aptamers were purified by standard desalting, while RNA oligonucleotides were purified by RNase-free high-performance liquid chromatography (HPLC). Two RNA oligonucleotides a 5-mer and a 10-mer sequences, randomly derived from the CHIKV genome and were used in the study (Supplementary Table S1). Additionally, a single-stranded DNA oligonucleotide (5′-CGTCGCTATA-3′) with the same sequence as RAC2 was purchased from Integrated DNA Technologies (Coralville, Iowa, USA) and included in the experiments. Double-stranded DNA was isolated from brain tissue of TgM83^+/−^ mice (Giasson et al., 2002) using the DNeasy Blood & Tissue Kit (Qiagen GmbH, Hilden Germany), which is optimized for genomic DNA extraction. The double-stranded DNA (dsDNA) was provided by Sara Reithoffer at the Institut für Physikalische Biologie, Heinrich-Heine-Universität Düsseldorf. A random single-stranded DNA (5′-TGACCATGGAGCCTGCCGTCTACTTCAAG-3′) was obtained from Sigma-Aldrich (St. Louis, MO, USA) and kindly provided by Dr. Jeannine Mohrlüder, from the Institut für Biologische Informationsprozesse, Strukturbiochemie (IBI-7), Forschungszentrum Jülich.

### 2.4 Enzymatic assay

To assess the activity of CHIKV nsP2^pro^ and the effects of nucleic acids, a FRET-based assay was performed using a synthesized fluorogenic peptide substrate, DABCYL-Arg-Ala-Gly-Gly-↓Tyr-Ile-Phe-Ser-EDANS (BACHEM, Bubendorf, Switzerland) (Eberle et al., 2021). The enzymatic assay was conducted in a 96-well plate with a final reaction volume of 100 µL per well. Unless otherwise stated, 1× PBS (pH 7.5) was used as the standard assay buffer condition.

Each well contained 10 µM nsP2^pro^, 9 µM fluorogenic substrate, and nucleic acids at a protease-to-nucleic acid molar ratio of either 1:1 (10 µM) or 1:3 (30 µM). Fluorescence intensities were measured (excitation at 340 nm, emission at 490 nm) every 30 seconds for 30 minutes at 37 °C.

To assess the influence of buffer conditions on nsP2^pro^ activity, a separate experiment was conducted using three different buffer environments: 20mM Phosphate buffer (pH 7.5), 20 mM Bis-Tris propane (pH 7.5), and 400 mM NaCl in Bis-Tris propane buffer (pH 7.5).

The activity and effects of nucleic acids on the protease were calculated using Eq. 1 (Eberle et al., 2021)

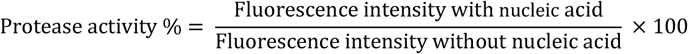

All assays were carried out in technical triplicates, and the resulting data were analyzed and presented as mean ± standard deviation (SD).

### 2.5 Structure predication of DNA aptamers

The secondary structures of DNA aptamers were predicted using the DINAMelt Server – Quikfold (https://www.unafold.org/quikfold.php) (Markham & Zuker, 2005). The Fast Folding – Energies & Structures algorithm was used to determine the most thermodynamically stable conformations. Predictions were carried out at 18 °C with two different Na^+^ concentrations: 137 mM and 10 mM. The sequence type was set to linear, and all other parameters were kept at their default values as provided by the server, including 5% suboptimality, a default window size, a maximum of 50 suboptimal foldings, and no limit on the distance between base-paired nucleotides.

### 2.6 CD spectroscopy

DNA and RNA samples were separately prepared in ddH_2_O and 1x PBS, respectively, to final concentrations of 30 µM for DNA aptamers and 50 µM for RNAs, enabling subsequent secondary structure analysis Circular Dichroism (CD) spectra were recorded at 18 °C using a 0.1 mm pathlength cuvette (Hellma GmbH & Co.KG, Muellheim, Germany), with a step size of 0.5 nm, over a wavelength range of 200–350 nm on a Jasco J-1100 spectropolarimeter (Jasco GmbH, Pfungstadt, Germany). Additionally, CD spectroscopy was employed to assess the conformational integrity of the purified nsP2^pro^ enzyme post-purification. The protease was diluted to a final concentration of 5 µM in a 10 mM sodium phosphate buffer (2.39 mM NaH_2_PO_4_, 7.6 mM Na_2_HPO_4_, pH 7.5). Measurements were conducted at 18°C using a 1 cm pathlength cuvette across the wavelength range of 190–260 nm, with seven replicate scans to ensure data reliability. For data processing, baseline spectra, averaged from multiple measurements, were subtracted from the corresponding sample spectra to yield corrected readings. The resulting data were expressed as molar ellipticity ([θ]), calculated using the Eq.2:

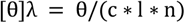

Where θ represents the ellipticity measured at the wavelength λ (in degrees), c is the protein concentration (mol/L), l is the cell path length (cm), and n is the number of residues in the protein.

### 2.7 *In silico* analysis

Models of CHIKV nsP2^pro^ and in complex with nucleic acids were generated using the AlphaFold 3 web server (Abramson et al., 2024). The structure of CHIKV nsP2^pro^ was retrieved from the PDB (code: 3TRK) and compared with the AlphaFold model (Supplementary Fig. S1).

To predict amino acids with the potential to interact with nucleic acids, the nsP2^pro^ alpha fold model was investigated with the PROBind webserver (https://www.csuligroup.com/PROBind/home) (Wu C. et al., 2025). A possible binding mode of different nucleic acids with CHIKV nsP2^pro^ was predicted by protein-nucleic acid blind docking at the web server HDOCK (Yan, Tao, He, & Huang, 2020), the results were compared with the models generated by the AlphaFold 3 web server (Abramson et al., 2024). Therefore, the AlphaFold modelled structures of CHIKV nsP2^pro^ and respective nucleic acids (ssDNA, DAC1, RAC1 and RAC2) were used for docking. The best HDOCK clusters were chosen based on the HDOCK score (Supplementary Fig. S2 and Table 2).

**Table 2.** Summary of chosen clusters after protein-nucleic acid blind docking using HDOCK.

|  | ssDNA | DAC1 | RAC2 | RAC1 |
| --- | --- | --- | --- | --- |
| **HDOCK score<sup>1</sup>** | -101.6 ± 7.3 | -109.2 ± 3.3 | -78.8 ± 3.9 | -71.5 ± 4.0 |
| **Cluster** | 2 | 1 | 1 | 7 |
| **Cluster size** | 17 | 96 | 20 | 13 |
| **RMSD<sup>2</sup>** | 1.0 ± 0.7 | 14.9 ± 0.4 | 0.6 ± 0.2 | 2.3 ± 0.1 |
| **VWD<sup>3</sup> energy** | -54.9 ± -5.3 | -45.5 ± 6.0 | -52.7 ± 1.4 | -36.9 ± 5.6 |
| **Electrostatic energy** | -323.8 ± 46.4 | -403.8 ± 35.7 | -222.4 ± 15.0 | -244.8 ± 30.2 |
| **Desolvation energy<sup>4</sup>** | 12.5 ± 0.8 | 14.8 ± 1.1 | 14.9 ± 1.1 | 13.4 ± 0.9 |
| **Restraints violation energy<sup>5</sup>** | 55.0 ± 18.7 | 22.7 ± 9.8 | 34.8 ± 23.1 | 19.5 ± 7.5 |
| **Z-score<sup>6</sup>** | -2.1 | -1.0 | -2.2 | -1.3 |

<sup>1</sup> Weighted sum of various energy terms, including van der Waals, electrostatic, and desolvation energies.
<sup>2</sup> RMSD from the overall lowest-energy structure.
<sup>3</sup> Van der Waals
<sup>4</sup> Energy penalty associated with bringing molecules into contact.
<sup>5</sup> Energy penalty assigned when a defined restraint (distance or dihedral angle), is not perfectly satisfied in a generated model.
<sup>6</sup> Indicates how many standard deviations a cluster's HADDOCK score deviates from the mean HADDOCK score of all clusters. A lower z-score (more negative) generally indicates a better, more favorable cluster.

All figures were created using the PyMOL Molecular Graphics System, Version 1.3, Schrödinger, LLC.

### 2.8 Statistical analysis

To assess the significance level (p-value), statistical analyses were performed using GraphPad Prism (version 5.0). One-way ANOVA followed by Tukey’s post hoc test was used to evaluate the differences between the groups and the control. Significance levels are indicated as follows: p < 0.05 (*), p < 0.01 (**), and p < 0.001 (***).

## 3. Results

### 3.1 DNA aptamer selection

Aptamers are short, single-stranded DNA or RNA molecules that fold into defined secondary and tertiary structures, allowing them to bind selectively and with high affinity to specific target molecules (Khan et al., 2022; Xu et al., 2023). To isolate DNA aptamers capable of binding the CHIKV nsP2^pro^, a High-Throughput Sequencing–Fluorescent Ligand Interaction Profiling (HiTS-FLIP) approach incorporating a primer-blocking strategy was employed. This method facilitated the direct selection of high-affinity aptamers from a large pool of randomized DNA sequences. Based on their strong binding affinity to the nsP2^pro^, ten candidate aptamers were selected for further analysis. The sequences and corresponding K_D_ values of the top ten aptamers, named DAC1–DAC10 (DNA against Chikungunya), are listed in Table 3, while the K_D_ fitting data are shown in Supplementary Fig. S3.

**Table 3.**
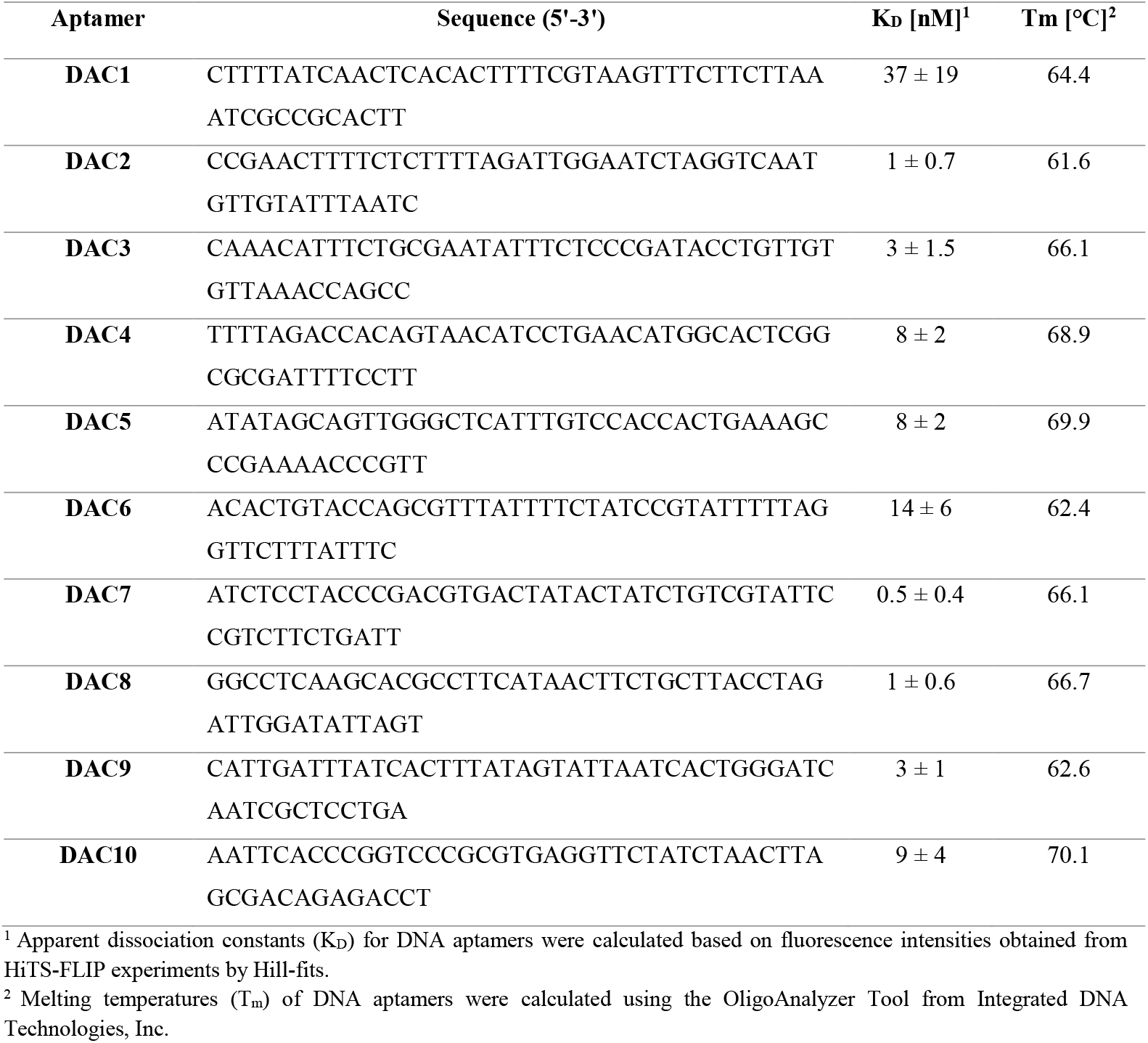
Sequences and properties of the selected DNA aptamers (DAC1–DAC10).

### 3.2 Secondary structure prediction of the nucleic acids

To evaluate the secondary structure of the selected DNA aptamers targeting CHIKV nsP2^pro^, the DINAMelt Server – Quikfold web tool was utilized (Markham & Zuker, 2005). This tool predicts folding conformations based on thermodynamic parameters (Supplementary Figs. S4-S5). The structural predictions revealed that all DNA aptamers could adopt secondary structures characterized by hairpin formations, although the lengths of the stems and sizes of the loops varied among the different sequences. To experimentally validate these computational predictions, CD spectroscopy was performed to assess the secondary structure content of the aptamers (Supplementary Figs. S6-S8). CD spectroscopy results showed similar structural features under both ddH_2_O and 1× PBS buffer conditions, with two positive peaks around 280 nm and 220 nm, and a negative peak near 245 nm. In addition to the DNA aptamers, two single-stranded RNA molecules RAC1 (5 nucleotides) and RAC2 (10 nucleotides), where “RAC” denotes RNA Against Chikungunya (see Supplementary Table S1), were analyzed by CD spectroscopy to evaluate their secondary structure characteristics. RAC1 exhibited a broad spectrum with low ellipticity, showing a weak positive peak around 270–280 nm, suggesting minimal or disordered secondary structure. In contrast, RAC2 displayed a more defined CD spectrum, with a clear positive peak at 270–280 nm and a distinct negative peak at around 240 nm, indicating the formation of a more ordered secondary structure (Supplementary Fig. S8).

### 3.3 Impact of DNA aptamers on CHIKV nsP2^pro^ enzymatic activity

To investigate the impact of nucleic acids on the enzymatic activity of nsP2^pro^, a primary activity assay was conducted in 1x PBS buffer using DNA aptamers targeting CHIKV nsP2^pro^ (DAC1–DAC10). The aptamers were mixed with 10 µM nsP2^pro^ at two molar ratios: 1:1 (10 µM) and 1:3 (30 µM). Fluorescence intensities (excitation at 340 nm, emission at 490 nm) were recorded every 30 seconds over a 30-minute. As shown in Fig. 1, all tested DNA aptamers significantly enhanced nsP2^pro^ activity. Among the ten aptamers variants, DAC1 exhibited the most pronounced effect, increasing enzymatic activity by approximately 600%. DAC2, DAC6, DAC8, and DAC9 induced up to 500% enhancement, while DAC3, DAC4, and DAC5 demonstrated moderate levels of activation (approximately 300-400%), suggesting differences in their ability to promote protease activity. The results also revealed that nsP2^pro^ activity was influenced by the aptamer-to-protease molar ratio. For most DAC variants, increasing the aptamer concentration from 10 µM to 30 µM did not further enhance activity. In fact, for DAC1, DAC2, DAC3, DAC4, and DAC6, higher concentrations led to a reduction in enzymatic activity, suggesting that higher aptamer concentrations do not lead to the higher activity rate in these cases. In contrast, DAC7, DAC8, DAC9, and DAC10 showed a slight increase in activity at the 1:3 ratio compared to 1:1, indicating only minimal additional enhancement.

**Fig. 1.**
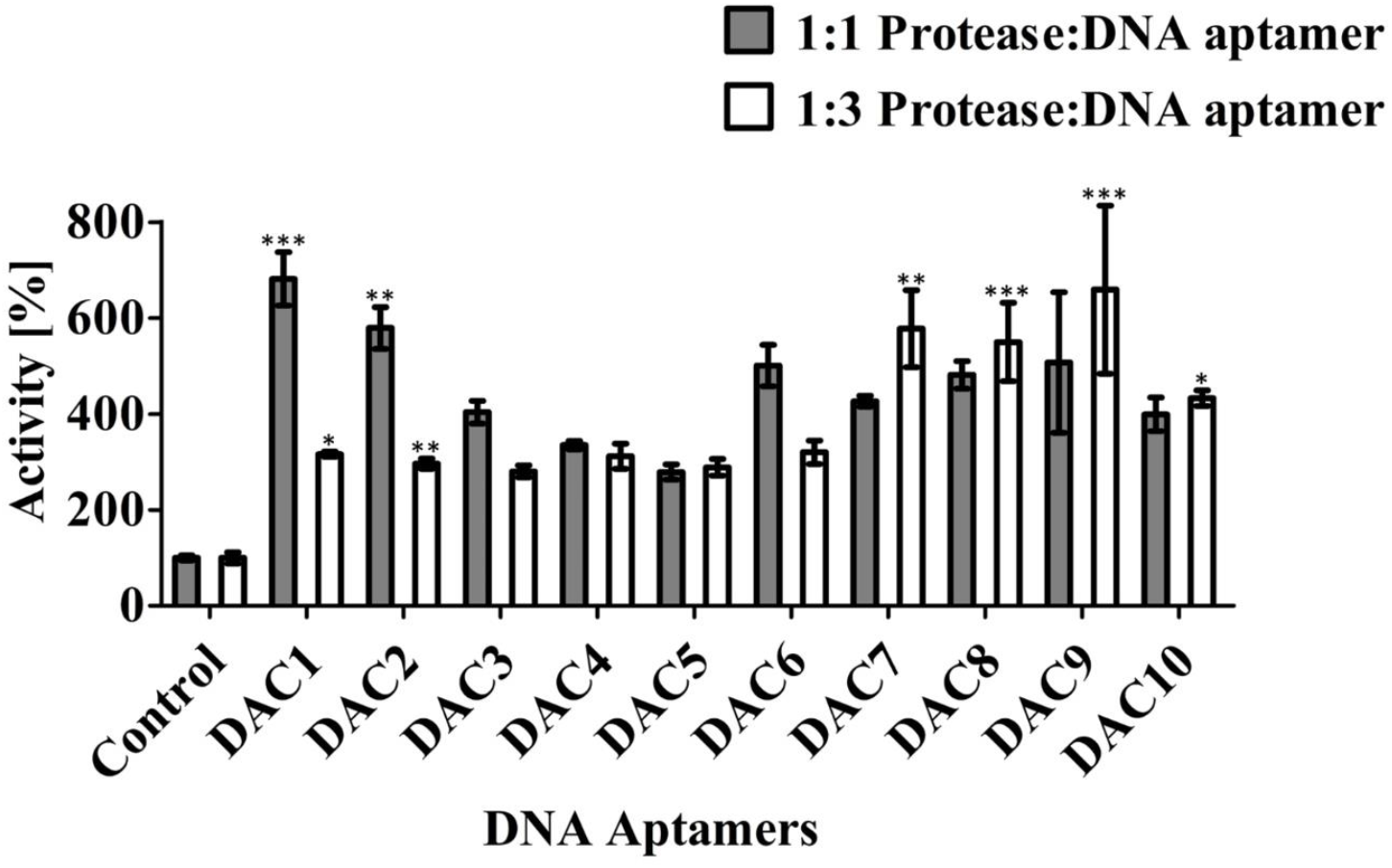
Enzymatic activity of nsP2^pro^ in the presence of DNA aptamers (DAC1–DAC10) at different molar ratios. primary activity assay of DNA aptamers was performed in 1× PBS at two different molar ratios: 1:1 and 1:3. The protease concentration was 10 µM, and the substrate concentration was maintained at 9 µM. The protease was mixed with DNA aptamers at 1:1 (10 µM, gray bars) and 1:3 (30 µM, white bars) molar ratios. Data from three independent experiments (n = 3) are presented as mean ± standard deviation (SD). Statistical significance was evaluated using one-way ANOVA followed by Tukey’s post hoc test. Asterisks indicate values that differ significantly from the respective control group. The significance levels are defined as follows: p < 0.05 (*), p < 0.01 (**) and p < 0.01 (***). If no asterisk is shown, the difference was not statistically significant (p > 0.05).

### 3.4 Impact of random ssDNA and dsDNA on the enzymatic activity of CHIKV nsP2^**pro**^

In Section 3.3, the impact of specific DNA aptamers on nsP2^pro^ activity was assessed. To determine the specificity of this effect, random single-stranded DNA (ssDNA) was used as a control. It was found that random ssDNA, at a 1:1 molar ratio with the protease, also enhanced protease activity, increasing it by up to 600% (Fig. 2A). To further investigate the effect of random DNA on nsP2^pro^ activity, the potential of random dsDNA, isolated from TgM83^+/−^ mice, was also examined. In contrast to ssDNA, dsDNA did not induce any detectable increase in nsP2^pro^ activity (Fig. 2A), indicating that the stimulatory effect is likely dependent on the single-stranded conformation of the nucleic acid.

**Fig. 2.**
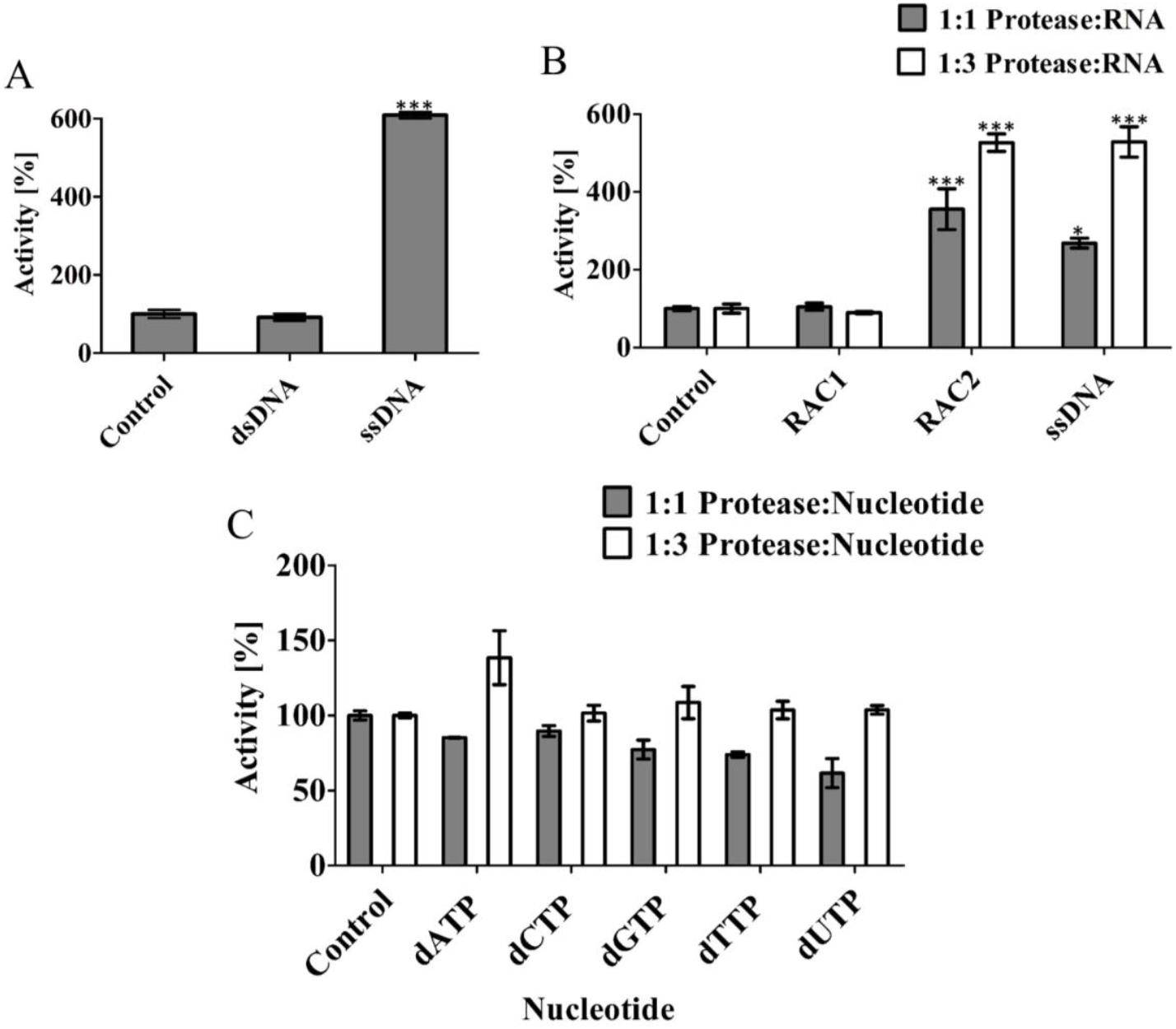
Influence of nucleic acids and nucleotides on nsP2^pro^ enzymatic activity. **(A)** Effect of random ssDNA and dsDNA on nsP2^pro^ activity. **(A)** Primary assay was conducted to assess the influence of random ssDNA (5′-TGACCATGGAGCCTGCCGTCTACTTCAAG-3′) and dsDNA on the enzymatic activity of nsP2^pro^. **(B)** Enzymatic activity of nsP2^pro^ in the presence of ssRNAs (RAC1–2) and ssDNA (5’-CGTCGCTATA-3’) corresponding to RAC2, tested at 1:1 (10 µM, gray bars) and 1:3 (30 µM, white bars) molar ratios. **(C)** Enzymatic activity of nsP2^pro^ in the presence of single nucleotides. Assays were performed in 1× PBS at two molar ratios: 1:1 (10 µM, gray bars) and 1:3 (30 µM, white bars). In all experiments, the protease concentration was 10 µM and the substrate concentration was maintained at 9 µM. Data are shown as mean ± standard deviation (SD) from three independent experiments (n = 3). Statistical significance was evaluated using one-way ANOVA followed by Tukey’s post hoc test. Asterisks indicate significant differences compared to the control group, with thresholds defined as: p < 0.05 (*), p < 0.01 (**), and p < 0.001 (***). If no asterisk is shown, the difference was not statistically significant (p > 0.05).

### 3.5 Impact of single-stranded RNA on CHIKV nsP2^pro^ enzymatic activity

Since CHIKV is an RNA virus that replicates in the host cell cytoplasm, it is of interest to evaluate whether RNA can influence the activity of nsP2^pro^. To investigate this, two ssRNA oligonucleotides, 5-mer and 10-mer sequences, randomly derived from the CHIKV genome were tested. These were designated RAC1 (5 nucleotides) and RAC2 (10 nucleotides) (Supplementary Table S1). The primary activity assay, conducted in 1x PBS buffer, showed that the shorter ssRNA, RAC1, did not enhance nsP2^pro^ enzymatic activity at either the 1:1 or 1:3 molar ratios (Fig. 2B) However, RAC2 significantly increased protease activity. At 10 µM, activity increased to around 300%, and at 30 µM, it rose to approximately 500% compared to the control (Fig. 2B). In addition, we tested a ssDNA oligonucleotide with the same sequence as RAC2. Interestingly, this ssDNA also enhanced nsP2^pro^ enzymatic activity, increasing it by approximately 260% at the 1:1 molar ratio and around 500% at the 1:3 molar ratio, similar to the effect observed with RAC2 (Fig. 2B).

### 3.6 Impact of nucleotides on the activity of CHIKV nsP2^**pro**^

To investigate whether individual nucleotides could enhance nsP2^pro^ activity, a primary activity assay was conducted under the same experimental conditions. Nucleotides were tested at concentrations of 10 µM and 30 µM. At the lower concentration (10 µM), no significant increase in protease activity was observed. However, at 30 µM, a modest increase in protease activity was detected (Fig. 2C). Specifically, dATP at 30 µM induced a modest increase of approximately 38%, which is substantially lower than the activity observed in presence of the ssDNA or ssRNA.

### 3.7 Impact of buffer composition on the activity of CHIKV nsP2^pro^ by DNA aptamer and ssRNA

To assess the effect of different buffers on nsP2^pro^ activity in the presence of nucleic acids, the ability of DAC8 (a representative DNA aptamer) and RAC2 (a 10-mer ssRNA) to enhance protease activity was evaluated under various buffer conditions. Assays were conducted using 10 µM nsP2^pro^ and two molar ratios of protease-to-nucleic acids: 1:1 and 1:3. Three buffer systems were tested: 20 mM Bis-Tris propane (pH 7.5), 20 mM phosphate buffer (pH 7.5), and 20 mM Bis-Tris propane supplemented with 400 mM NaCl (pH 7.5). As shown in Fig. 3A–B, both DAC8 and RAC2 enhanced protease activity in Bis-Tris propane and phosphate buffers at both molar ratios. Notably, DAC8 induced a more pronounced increase in activity than RAC2 at the 1:1 ratio. However, at the 1:3 ratio, RAC2 exhibited greater activation than DAC8 in Bis-Tris propane buffer, while in phosphate buffer, both nucleic acids produced similar effects. In contrast, under high-salt conditions (400 mM NaCl), neither DAC8 nor RAC2 was able to stimulate nsP2^pro^ activity at either molar ratio. These findings suggest that elevated ionic strength inhibits nucleic acid-mediated enhancement of protease activity and underscore the critical role of buffer composition in modulating nsP2^pro^ enzymatic function.

**Fig. 3.**
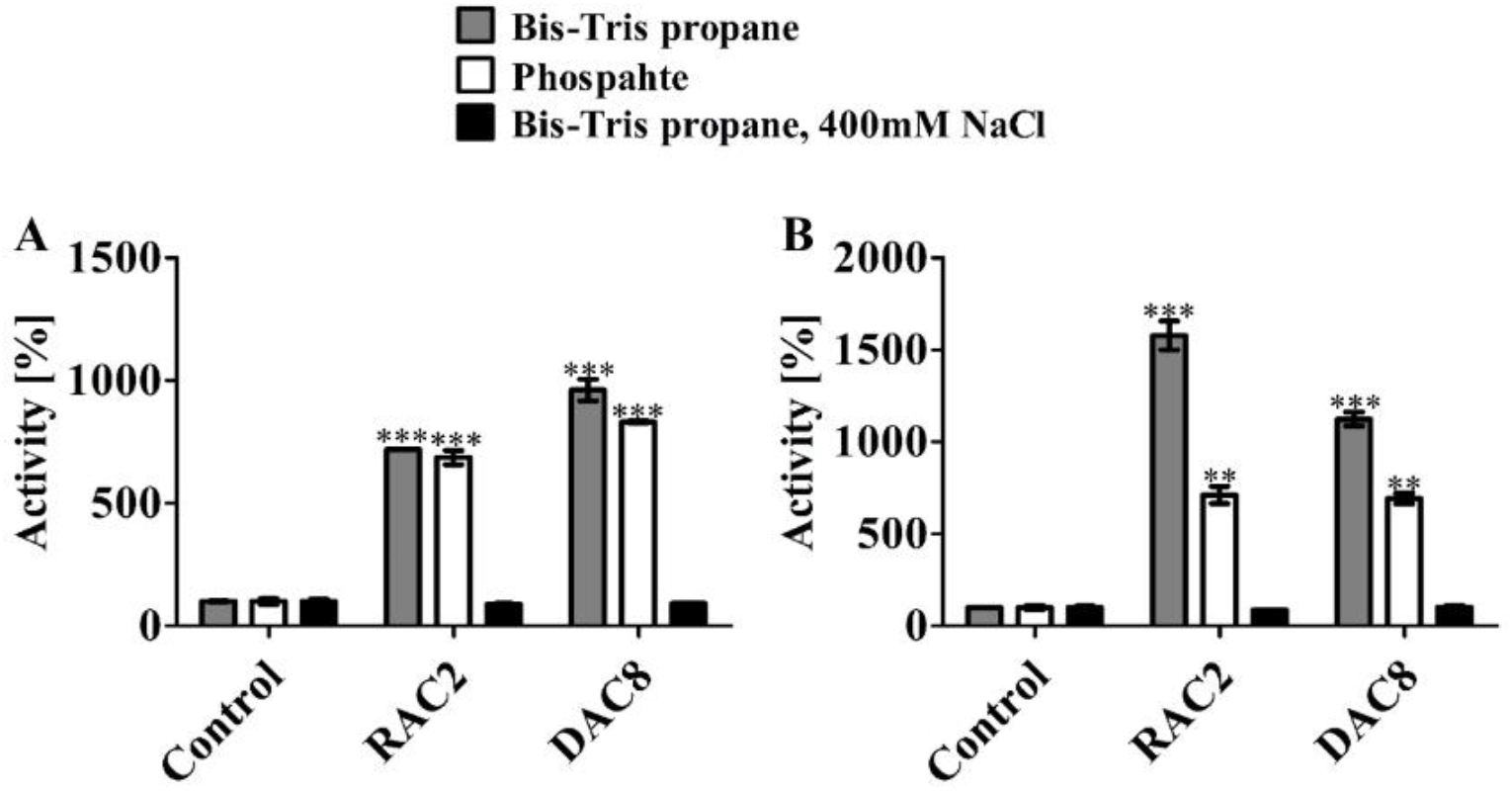
Effect of buffer composition on the activity of nsP2^pro^ by nucleic acids. The activity of nsP2^pro^ by DAC8 (a DNA aptamer) and RAC2 (ssRNA) was assessed in the presence of three different buffer conditions: (1) 20 mM Bis-Tris propane, pH 7.5, (2) 20 mM phosphate buffer, pH 7.5, and (3) 20 mM Bis-Tris propane, 400 mM NaCl, pH 7.5. Nucleic acids were tested at molar ratios of 1:1 **(A)** and 1:3 **(B)** with nsP2^pro^ (10 µM). The substrate concentration was kept at 9 µM. Data are presented as the mean ± SD from three independent experiments (n = 3). Statistical significance was evaluated using one-way ANOVA followed by Tukey’s post hoc test. Asterisks indicate significant differences compared to the control group, with thresholds defined as: p < 0.05 (*), p < 0.01 (**), and p < 0.001 (***). If no asterisk is shown, the difference was not statistically significant (p > 0.05).

### 3.8 Possible binding mode of nucleic acids with CHIKV nsP2^**pro**^

CHIKV nsP2^pro^ consists of an N-terminal papain-like protease domain, directly followed by a Ftsj MTase-like domain (Fig. 4A). The surface of the MTase-like domain possesses clusters of positive charges, located near the protease active site loop carrying His1083 (Fig. 4B), which may propagate the binding of the negative charged phosphate backbone of nucleic acids.

**Fig. 4.**
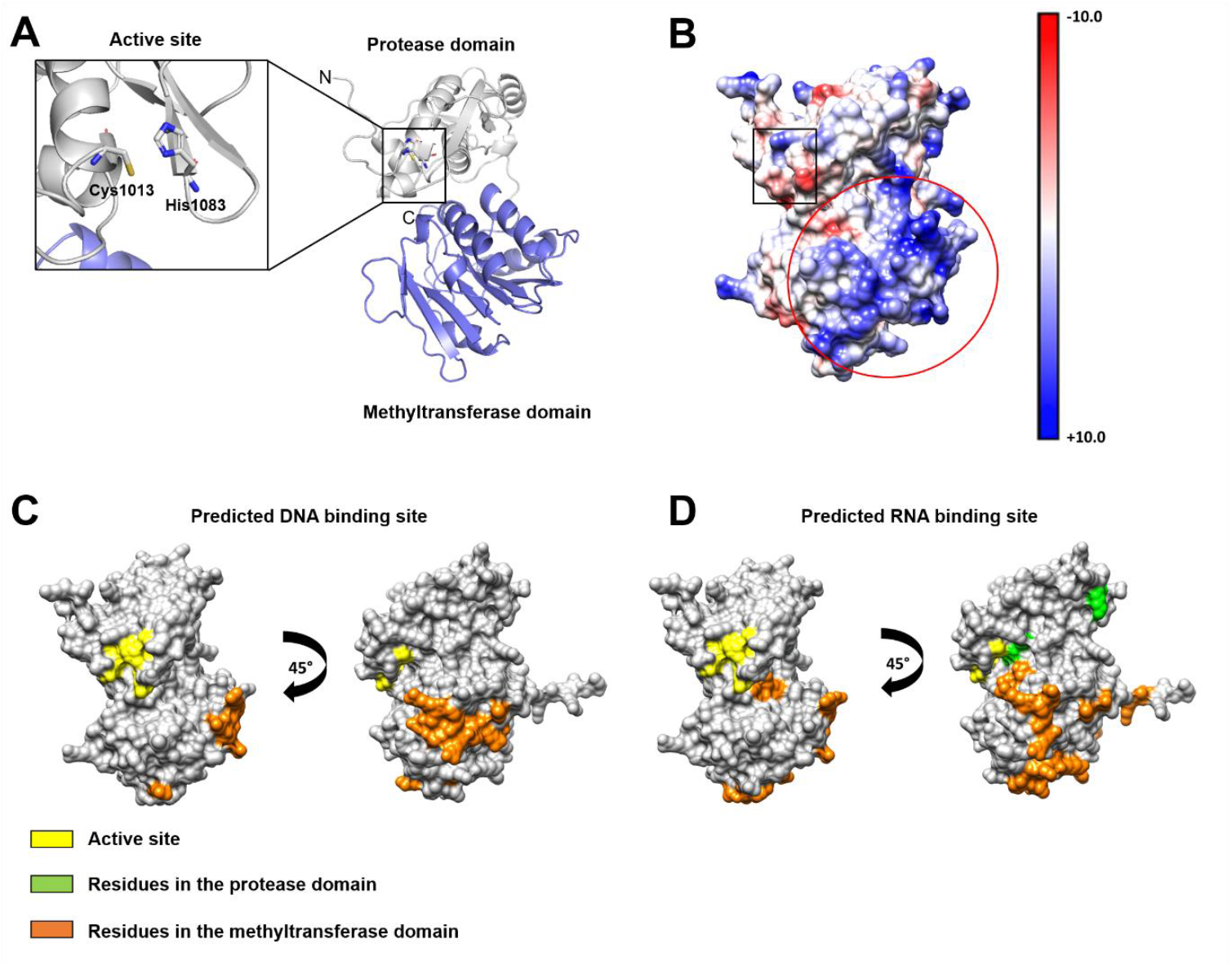
Investigation of the nsP2^pro^ AlphaFold model with regard to possible nucleic acid binding regions. A structural comparison between the crystal structure of the protein (PDB code: 3TRK) and the generated AlphaFold model is shown in supplementary Fig. S1. **(A)** nsP2^pro^ is shown in ribbon view, the papain-like cysteine protease is colored in grey and the Ftsj methyltransferase (MTase)-like domain in blue. The black box labels the position of the active site residues Cys1013 and His1083. **(B)** Coulombic surface of nsP2^pro^, the red circle labels a strongly positively charged area at the MTase domain, the black box labels the position of the active site residues. The online tool PROBind was used to predict amino acids that can take part in interactions with DNA and RNA. The protein is shown in surface view and the predicted amino acids are labeled in orange (MTase domain) and green (Protease domain). The position of the active site is labeled in yellow. **(C)** Predicted DNA binding site. **(D)** Predicted RNA binding site.

The online tool PROBind was used to predict amino acids that are involved in DNA and RNA binding (Fig. 4C-D). Thereby, 14 residues were predicted to interact with DNA, located mainly at a surface cluster in the MTase domain (Fig. 4C). 23 residues were predicted to interact with RNA, where 16 residues are located at the MTase domain and 7 residues at the active site loop carrying His1083 on the protease domain. Some areas seem to overlap regarding the DNA and RNA interaction, the predicted amino acids are shown in the nsP2^pro^ sequence at supplementary Fig. S9.

To predict a possible binding area, models of complexes between CHIKV nsP2^pro^ and different nucleic acids were generated using AlphaFold 3. The tested nucleic acids included ssDNA (5′-CGTCGCTATA-3’), DAC1 (5′-CTTTTATCAACTCACACTTTTCGTAAGTTTCTTCTTAAATCGCCGCACTT-3`), RAC1 (5′-CGUCG-3′) and RAC2 (5′-CGUCGCUAUA-3′). Additionally, blind docking was performed using HDOCK (Yan et al., 2020). The positions of the nucleic acids in the complexes with the protein were compared between both approaches, as shown in Fig. 5. To demonstrate a stable pose within the different clusters, an overlay of all predicted docking poses is shown in supplementary Fig S10. The results of both approaches demonstrated that the tested nucleic acids were docked without exception to the MTase domain. Interestingly, RAC2 showed almost the same pose between both approaches (Fig. 4D). These observations let us assume that the nucleic acids mainly interact with the MTase domain.

**Fig. 5.**
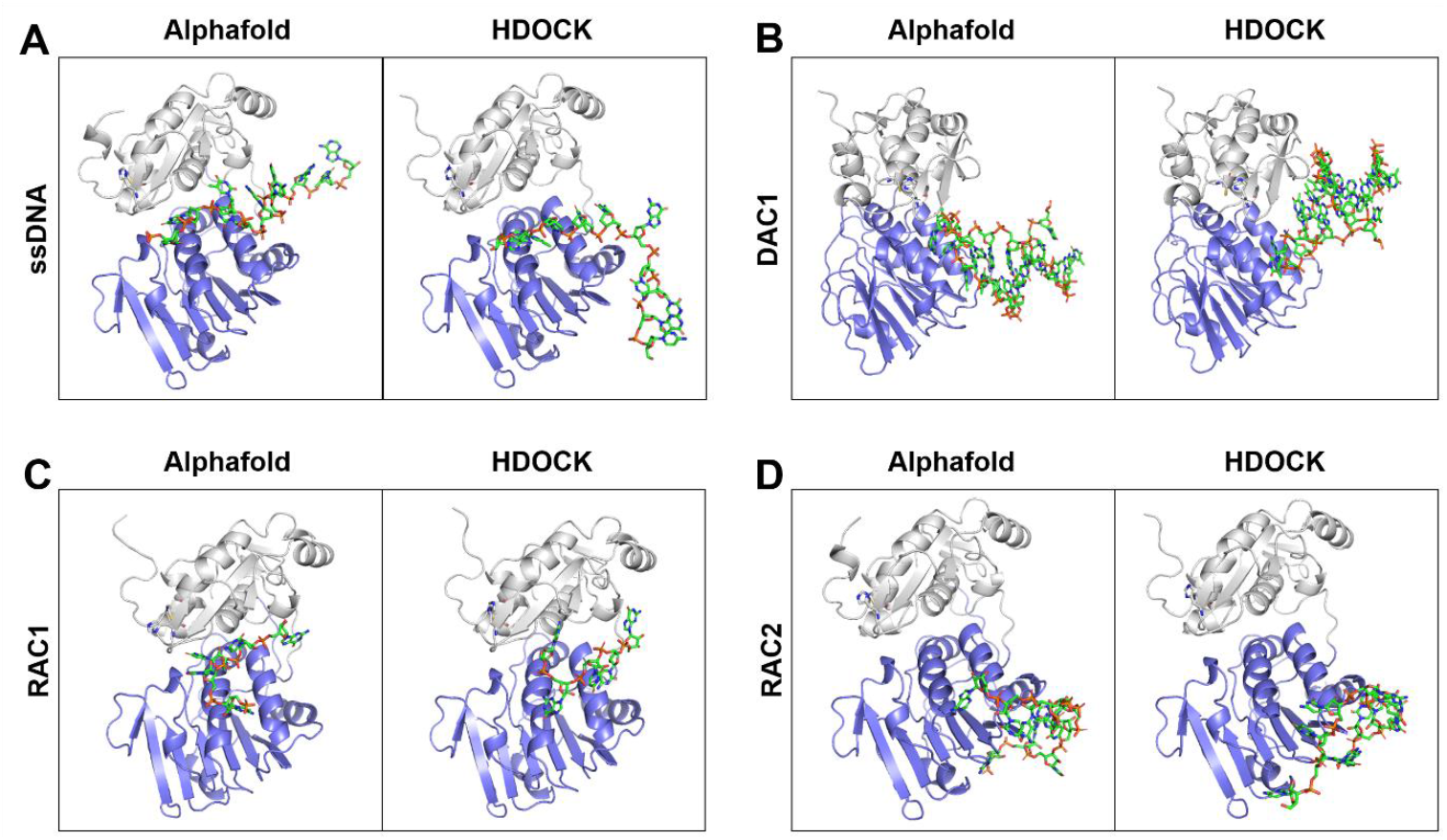
Binding pose of the different nucleic acids with nsP2^pro^ after complex generation using AlphaFold and docking using HDOCK. The target protein is shown in ribbon view, the papain-like cysteine protease is colored in grey and the Ftsj methyltransferase (MTase)-like domain in blue. The nucleic acids are shown as green sticks. One representative AlphaFold model is shown, the overlay of the five generated models is presented in supplementary Fig. S10 and show especially in the binding region a stable pose. A representative structure of the best-scoring cluster following protein-nucleic acid blind docking using HDOCK is also shown.

## 4. Discussion

The Chikungunya virus (CHIKV) is a rapidly emerging arthropod-borne pathogen whose increasing global distribution is related to climate and environmental changes. Central to CHIKV replication is its nonstructural protein 2 (nsP2), a multifunctional enzyme that plays an important role as a viral protease (nsP2^pro^) by mediating the cleavage of the viral polyprotein. Several studies have reported that interactions between viral proteins and nucleic acids are pivotal for mediating key aspects of the viral life cycle. For example, the Rep protein of adeno-associated virus type 2 binds single-stranded DNA (ssDNA), a process that facilitates viral replication (Stracker et al., 2004) and HIV-1 protease has been shown to bind viral RNA, potentially augmenting its catalytic activity (Potempa et al., 2015). In other cases, the binding of viral RNA to proteases such as those of the Foamy virus (Hartl et al., 2011) and the 3C^pro^ protease of poliovirus (Campagnola & Peersen, 2023) has been reported to activate protease function. Not only RNA but also DNA can mediate protease activation. For instance, host DNA has been shown to activate the 3C^pro^ of Seneca Valley virus (L. Wu et al., 2025), highlighting the potential role of host nucleic acids in modulating viral protein activity. Interestingly, nucleic acid interactions do not always facilitate viral replication; in some cases, they can inhibit it. For example, RNA aptamers have been identified that bind to the NS3 protease of hepatitis C virus, inhibiting its function in polyprotein processing (Urvil et al., 1997). Similarly, RNA aptamers targeting HIV-1 PR have been reported to block its enzymatic activity (Duclair et al., 2015). In light of these findings, this study focused on CHIKV nsP2^pro^ to assess the impact of nucleic acid interactions on its protease activity. HiTS-FLIP was employed to identify ssDNA aptamers that bind to nsP2^pro^. Ten aptamers (designated DAC1– DAC10) were selected based on their binding affinity (Table 3) and were subsequently tested for their effect on protease activity. To evaluate the effect of these aptamers on nsP2^pro^ activity, primary activity assays were conducted. The results showed that all ssDNA aptamers had the potential to enhance the activity of the target protease (Fig. 1). Among the ten DAC variants, DAC1 exhibited the most pronounced effect, increasing enzymatic activity by 600% (Fig. 1). DAC2, DAC6, DAC8, and DAC9 also induced substantial increases, reaching up to 500%. In contrast, DAC3, DAC4, and DAC5 showed moderate enhancement at a 1:1 ratio relative to the protease concentration (10 µM). However, when the DNA concentration was increased to a 1:3 ratio, a reduction in protease activity was observed, with DAC1-induced activity decreasing to approximately 300% compared to the 1:1 condition. This reduction may be attributed to the elevated ssDNA concentration, which could promote protein precipitation and aggregation-a phenomenon reported for proteins such as tau and α-synuclein, where nucleic acids have been shown to induce aggregation (Yin, Chen, & Liu, 2009). To ensure that the observed activity increase was due to protease–nucleic acid interactions and not simply optical interference from the aptamers themselves, control experiments were conducted in the absence of substrate. Reactions were prepared with buffer alone, protease plus buffer, and protease with buffer and DNA aptamers, excluding the fluorogenic substrate. No detectable absorbance was observed in the lack of substrate, confirming that the increased signal in the complete assays was attributable to enzymatic activity, not background fluorescence or absorbance changes caused by the aptamers (Supplementary Fig. S11). To determine whether the observed increase in nsP2^pro^ activity was due to specific binding by the selected ssDNA aptamers or could also result from non-specific interactions, a random ssDNA sequence was tested. Interestingly, as shown in Fig. 2A, the random ssDNA also enhanced protease activity, suggesting that the effect is not sequence-specific. To further explore this phenomenon, the effect of a random dsDNA, isolated from TgM83^+/−^ mice, was examined. In contrast to ssDNA, the random dsDNA did not cause any enhancement in protease activity compared to the control (Fig. 2A), suggesting that the stimulatory effect is dependent on the single-stranded nature of the nucleic acid. Since CHIKV is a ssRNA virus that replicates in the cytoplasm, it is likely that nsP2^pro^ naturally encounters ssRNA during infection. To evaluate whether ssRNA could similarly influence protease activity, two random ssRNA oligonucleotides were tested: a 5-mer (RAC1) and a 10-mer (RAC2), both derived from the CHIKV genome (Table S1). Activity assays revealed that RAC1 did not enhance nsP2^pro^ activity, whereas RAC2 significantly increased activity, reaching approximately 600% (Fig. 2B). These findings suggest that ssRNA, like ssDNA, can enhance protease activity, but the effect is length-dependent and requires a minimum RNA length. To confirm the length dependency, an additional assay was performed using single nucleotides. At a 1:1 molar ratio with the protease, no change in activity was observed; however, at a 1:3 ratio, a slight increase was detected (Fig. 2C). This result aligns with the ssRNA experiments and further supports the conclusion that enhancement of nsP2^pro^ activity by nucleic acids requires a minimal oligonucleotide length.

The influence of buffer composition on nucleic acid-mediated activation of nsP2^pro^ was systematically evaluated using two representative oligonucleotides: DAC8 (ssDNA aptamer) and RAC2 (ssRNA). Three buffer systems were tested: 20 mM Bis-Tris propane (pH 7.5), 20 mM phosphate buffer (pH 7.5), and 20 mM Bis-Tris propane supplemented with 400 mM NaCl (pH 7.5). As shown in Fig. 3A–B, both DAC8 and RAC2 significantly enhanced protease activity in Bis-Tris propane and phosphate buffers without added salt. At the 1:1 ratio, DAC8 induced a stronger activation compared to RAC2, while at the 1:3 ratio, RAC2 exhibited greater enhancement in Bis-Tris propane. In phosphate buffer, both oligonucleotides led to similar effects. However, in the presence of high salt (400 mM NaCl), the activation was completely abolished for both DAC8 and RAC2, irrespective of the molar ratio. These results indicate that the stimulatory interaction between nucleic acids and nsP2^pro^ is highly sensitive to ionic strength. The loss of activity in high-salt conditions suggests that electrostatic interactions play a critical role in facilitating protease activity. This observation aligns with findings from other studies, such as those involving the hsRosR protein, demonstrated that high salt concentrations can disrupt protein-nucleic acid binding by altering solvent properties or changing the electrostatic environment of the protein and DNA and the correlation in between aptamer affinity and pI value of target proteins (Kutnowski et al., 2019; Drees et al., 2023). Therefore, the effect of high salt on nsP2^pro^ activity likely stems from the disruption of protein-nucleic acids interactions, emphasizing the importance of buffer conditions for nucleic acid-mediated protease activation. To gain a better understanding of the structural basis of the DNA aptamers, the secondary structures of the selected DNA aptamers were predicted using the DINAMelt Server – Quikfold web tool. (Markham & Zuker, 2005). The predictions revealed that all sequences adopted secondary structures characterized by hairpin formations, although the lengths of the stems and the sizes of the loops varied among the different aptamers (Supplementary Figs. S4–S5). CD spectroscopy further confirmed the presence of these structured forms in both ddH_2_O and 1× PBS conditions. The spectra of all DNA aptamers exhibited two distinct positive peaks around 280 nm and 220 nm, along with a strong negative peak near 245 nm. A pattern consistent with the B-form DNA conformation (Kypr, Kejnovská, Renčiuk, & Vorlíčková, 2009) (Supplementary Figs. S6–S8). These spectral features corroborate the computational predictions and indicate that all DNA aptamers retain stable secondary structures in solution. Furthermore, comparative CD analysis of these DNA aptamers with two ssRNA molecules (RAC1 and RAC2) revealed marked structural differences. RAC2 exhibited a spectral profile characteristic of the A-form RNA helix, including a moderate positive band near 270 nm, reflecting a well-structured RNA fold (Sosnick, Fang, & Shelton, 2000). In contrast, RAC1 showed significantly diminished ellipticity across the spectrum, consistent with an unfolded or denatured state. This interpretation is in agreement with previous findings, where unfolded RNAs lack the defined CD spectral features associated with stable secondary structures (Sosnick et al., 2000) (Supplementary Fig. S8). These findings suggest that the secondary structure of nucleic acids may influence their binding capacity to nsP2^pro^ and contribute to their ability to enhance enzymatic activity.

Finally, nsP2^pro^ was investigated to specify possible nucleic acid binding regions at the protein surface, using the PROBind webserver (Wu C. et al., 2025). Predicted residues involved in DNA and RNA interaction differ in quantity and the localization in the protein. 14 amino acid residues were predicted to interact with DNA and 23 residues with RNA. DNA binding regions are exclusively localized at the MTase domain, also the majority of the RNA binding residues, but the protease domain carries five amino acids with the potential to interact with RNA. The proposed residues involved in DNA interaction included two main regions _1281_RSSR_1284_ and _1305_DNGRR_1309_. Contrary, predicted regions for RNA interaction included _1308_RRN_1310, 1241_QML_1243, 1183_TKR_1185, 1165_K(I)NGH_1169_ (predicted without the I) and five residues in the protease domain, _1054_NE_1055, 1050_E and _47_YS_48_ (Supplementary Fig. S9).

Viral proteins utilize various nucleic acid binding motifs, including the RNA recognition motif (RRM) and the zinc finger motif (Frequently involved in DNA binding), to interact with viral and host nucleic acids. Moreover, arginine plays an important role in several RNA binding motifs in virus proteins, e.g. arginine rich motif (ARM) (Casu, Duggan, & Hennig, 2013), HR motif (Ogino & Banerjee, 2010), RG/RGG motif (Iacovides, O’Shea, Oses-Prieto, Burlingame, & McCormick, 2007), SR/RS motif (Nikolakaki & Giannakouros, 2020). Arginine has a high occurrence in the predicted nucleic acid interaction regions of nsP2^pro^, these regions differ slightly to the typical motifs mentioned above. Viral proteins often exhibit significant variation in their RNA and DNA binding motifs. This variation arises from evolutionary pressures and the need for viruses to efficiently interact with the host cell’s RNA machinery and evade the host’s immune system (Garcia-Moreno et al., 2019). However, arginine plays a crucial role in nucleic acid interactions, particularly in RNA, due to its unique ability to form strong, multivalent interactions. Arginine can interact with the phosphate backbone and base pairs in RNA, forming salt bridges and hydrogen bonds that are essential for recognition and binding (B. Martin et al., 2022).

To predict a possible nucleic acid binding region on nsP2^pro^, molecular docking of four different nucleic acids, ssDNA, DAC1, RAC1 and RAC2 with the protein was performed using AlphaFold 3 and HDOCK. During the docking experiments no amino acids or regions were predefined, where the nucleic acids supposed to bind, we performed blind docking, with two different approaches, AlphaFold 3 and HDOCK. The results indicated that the different nucleic acids interact in the regions predicted by PROBind. Comparison of the nucleic acid positions between the Alphafold 3 models demonstrated that there were small differences between the binding poses of the tested nucleic acids between the AlphaFold 3 and HDOCK approach. However, our results let us assume, that the nucleic acids interact with the nsP2^pro^ MTase domain, but to generate valid information about the nucleic acid binding region of nsP2^pro^ further experiments are necessary.

Interestingly, a previous study from our group demonstrated an interdomain motion leading to closed and open conformations of the nsP2^pro^ active site. Thereby, a flap formed by residues A1080 to H1083 can adopt positions either close to or far from the residues _1202_NLELGL_1207_ in the MTase domain. When the active site adopts a more open conformation, the flap moves farther away from the loop on the C-terminal domain. (Mastalipour et al. 2025). Nearby the MTase _1202_NLELGL_1207_ loop are predicted nucleic binding regions are located. Especially, _1241_QML_1243_ is in direct neighborhood of this loop, 1242M and L1243 have a distance of 3.4 to 3.5 Å to the _1202_NLELGL_1207_ loop, which makes the formation of van der waals (VDW) interactions between both regions highly possible (Supplementary Figs S12 and S13). VDW interactions occur at distances typically between 2.5 and 4.6 angstroms (Å), averaging around 3.6 Å (Qi, Wang, & Ren, 2016). We assume that nucleic acids may interact with _1241_QML_1243_ and disturb the interaction to _1202_NLELGL_1207_, with the result that the active site exists in the open conformation. Moreover, the residues _1054_ NE_1055, 1050_E and _47_YS_48_ are located at a helix beside the active site. Nucleic acid binding in this region may also induce a conformational change that results in the open conformation of the protease active site.

## Conclusion and Future Perspectives

In this study, it was demonstrated that single-stranded nucleic acids-including both DNA aptamers and structured RNA fragments-can significantly enhance the catalytic activity of the Chikungunya virus nsP2^pro^. This enhancement appears to be influenced by several factors: the single-stranded nature of the nucleic acid, a minimum oligonucleotide length, the presence of stable secondary structures, and sensitivity to ionic strength. Molecular docking analyses predicted that these nucleic acids interact with the methyltransferase domain of nsP2^pro^; however, this binding site remains to be experimentally validated. Further studies are needed to confirm the precise binding interface and determine whether this interaction occurs under physiological conditions. Future investigations should focus on assessing the *in vivo* relevance of these interactions during CHIKV infection and replication, as well as on high-resolution structural studies to define the molecular details of the nucleic acid–nsP2^pro^ complex.

## CRediT authorship contribution statement

**Mohammadamin Mastalipour:** Conceptualization, Methodology, Formal analysis, Data curation, Writing – original draft, Writing – review & editing. **Mônika Aparecida Coronado:** Formal analysis, Writing – review & editing. **Ruth Anasthasia Siahaan:** Investigation. **Alissa Drees:** Methodology, Investigation, Data curation, Writing – review & editing. **Christian Ahlers:** Software, Investigation, Analysis, Writing – review & editing. **Markus Fischer:** Resources, Supervision, Writing – review & editing. **Dieter Willbold:** Resources, Writing – review & editing. **Raphael Josef Eberle:** Conceptualization, Supervision, Writing – review & editing. All authors have read and agreed to the published version of the manuscript.

## Declaration of generative AI and AI-assisted technologies in the writing process

ChatGPT (OpenAI) was utilized to assist with grammar and spelling checks to enhance the readability of the manuscript. All content was subsequently reviewed and edited by the authors, who take full responsibility for the final version of the text. In addition, selected figures were created using the BioRender platform.

## Acknowledgements

This work was supported by funding and resources from the Jürgen Manchot Stiftung, whose contribution was essential to the advancement of this research

