## Supplementry and material for "Single-Stranded nucleic acid binding enhances the *in vitro* catalytic activity of Chikungunya virus nsP2 protease"

**Table of content**

Supplementary figure S1. Structural comparison of the nsP2$^{pro}$ structure and the generated nsP2$^{pro}$ AlphaFold 3 model.

Supplementary figure S2. Van der Waals energy, electrostatic energy and restraints energy of the generated clusters after protein-nucleic acid blind docking using HDOCK.

Supplementary figure S3. $K_D$ fitting of DNA aptamers DAC1–DAC10.

Supplementary figure S4. Predicted secondary Structures of DAC1–DAC5. Panels.

Supplementary figure S5. Predicted secondary Structures of DAC6–DAC10.

Supplementary figure S6. Secondary structure of DAC1–DAC4.

Supplementary figure S7. Secondary structure of DAC5–DAC8.

Supplementary figure S8. Secondary structure of DAC9–DAC10 and RAC1-RAC2.

Supplementary figure S9. Amino acids of predicted nucleic acid binding regions in the nsP2$^{pro}$ sequence.

Supplementary figure S10. Overlay of the predicted nucleic acid positions for AlphaFold models and HDOCK results.

Supplementary figure S11. Control experiments confirm substrate-specific fluorescence signal in the nsP2$^{pro}$ activity assay

Supplementary Table S1 Sequences and properties of the RNAs used in this study

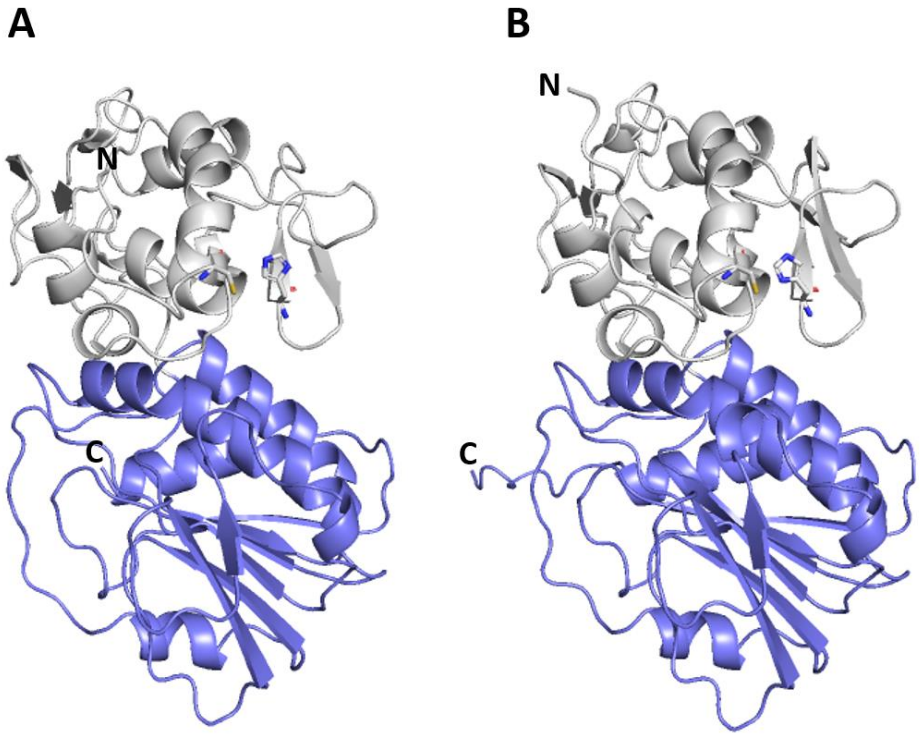


**Fig. S1. Structural comparison of the nsP2$^{pro}$ structure and the generated nsP2$^{pro}$ AlphaFold model.** Both structures are shown in ribbon view, the papain-like cysteine protease is colored in grey and the Ftsj methyltransferase (MTase)-like domain in blue. An overlay of both structures indicated a RMSD value of 0.435 (2165 to 2165 atoms). **(A)** nsP2$^{pro}$ crystal structure (PDB code: 3TRK) and **(B)** nsP2$^{pro}$ AlphaFold 3 model.

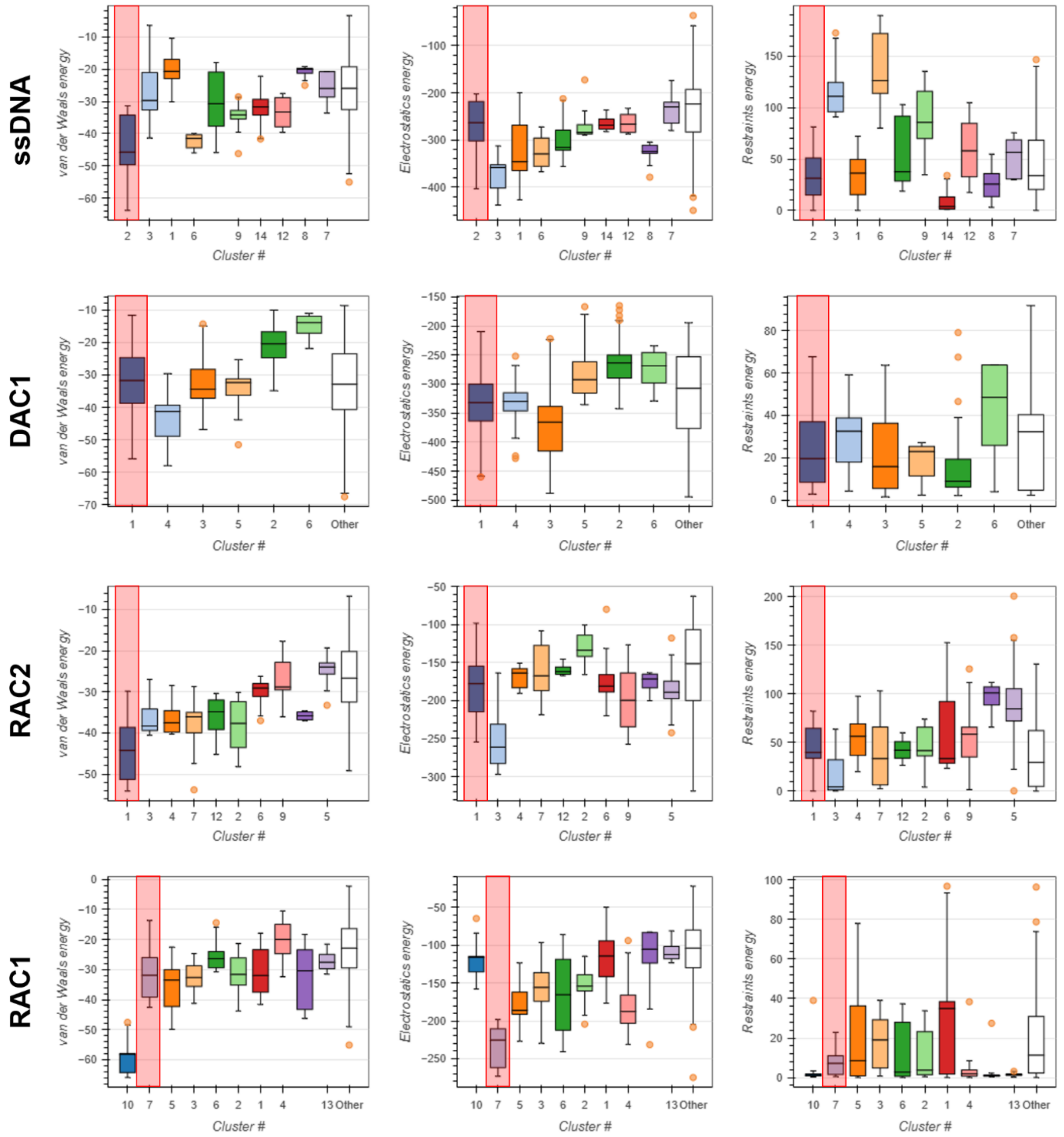


**Fig. S2. Van der Waals energy, electrostatic energy and restraints energy of the generated clusters after protein-nucleic acid blind docking using HDOCK.** Red boxes label the representative cluster for each nucleic acid.

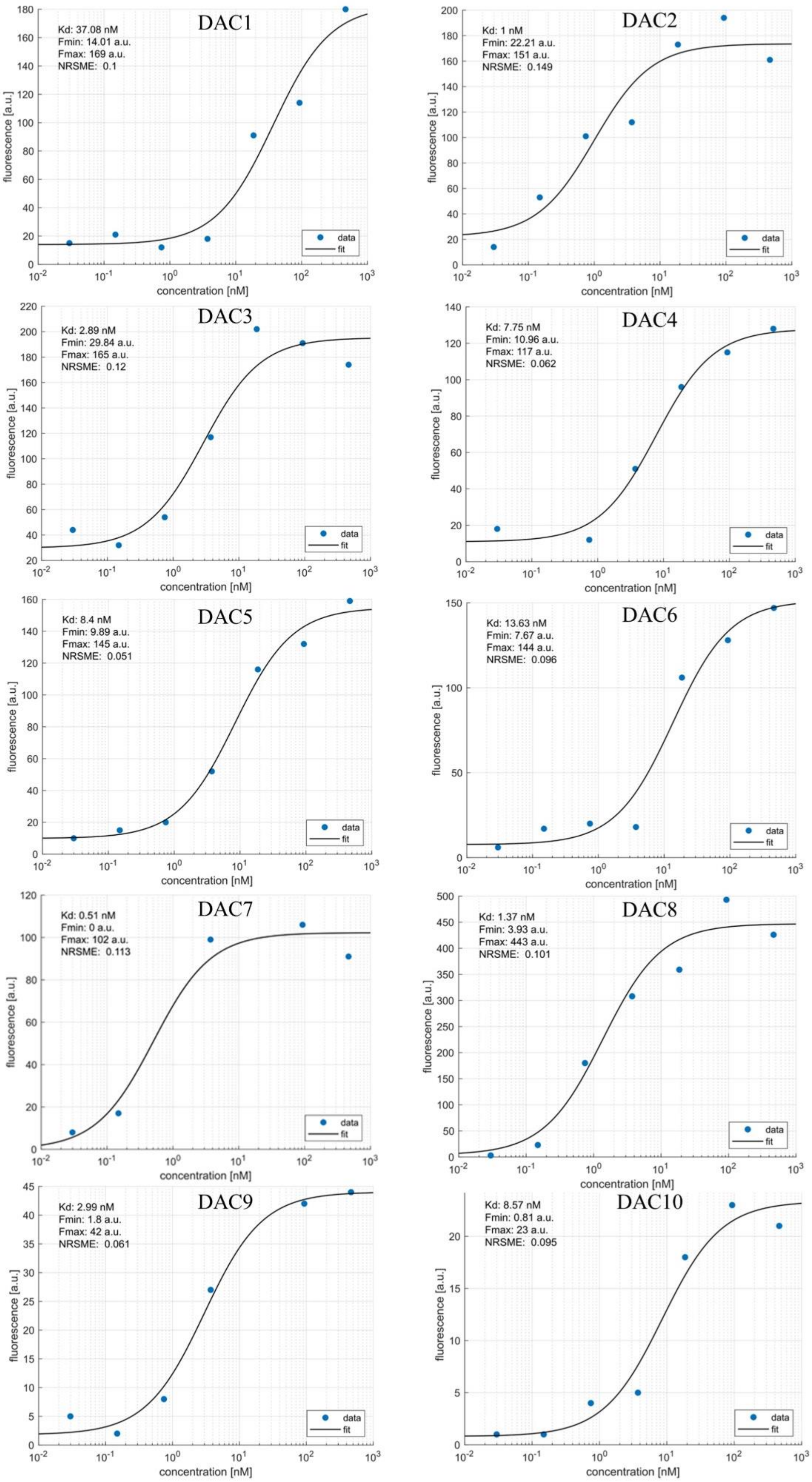


**Fig. S3. $K_D$ fitting of DNA aptamers DAC1–DAC10**. Binding affinities for DAC1–DAC10 were determined using HiTS-FLIP, and the data were fitted using a nonlinear Hill-fit.

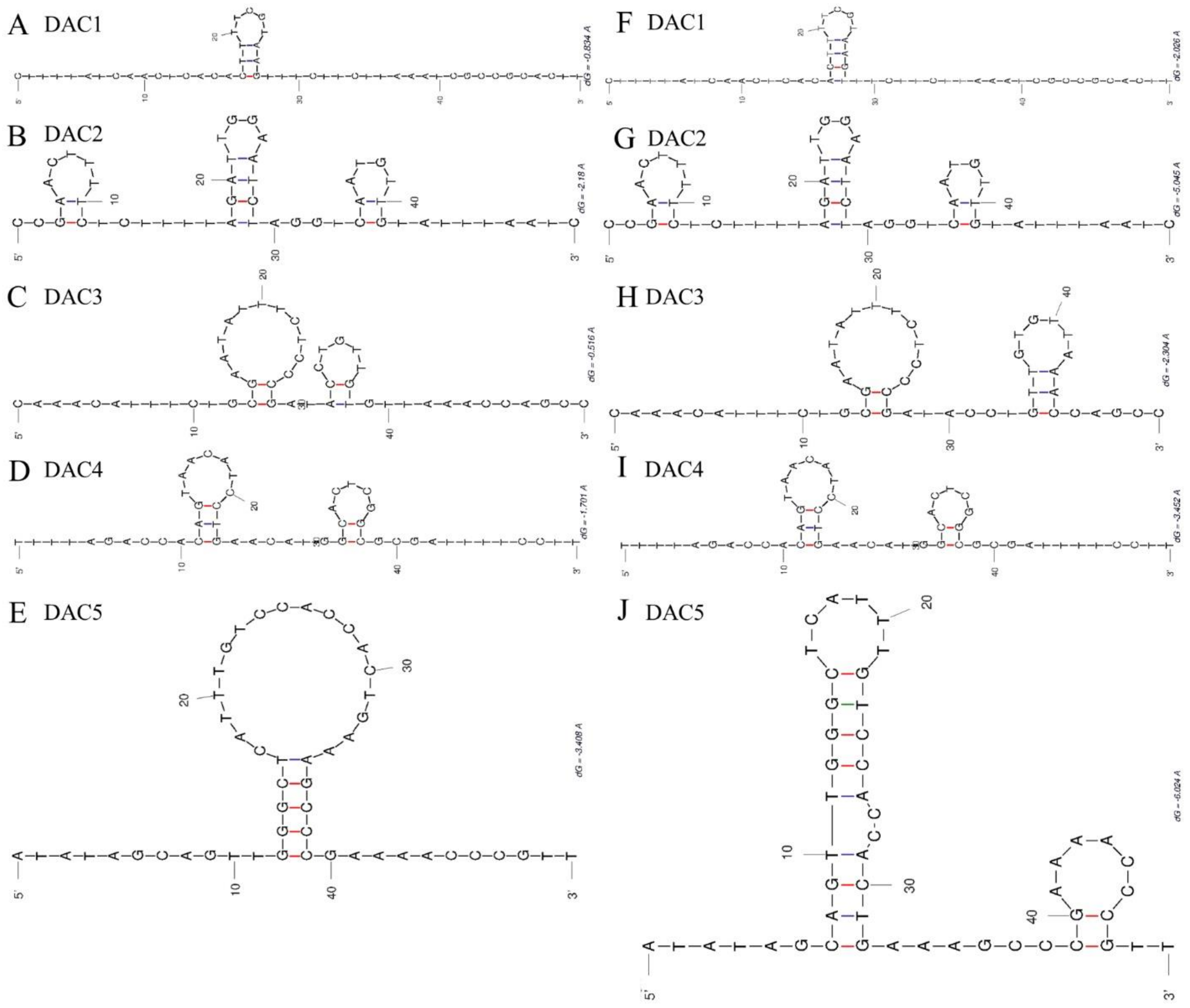


**Fig. S4. Predicted secondary structures of DAC1–DAC5.** Panels (A–E) show the predicted secondary structures of DAC1–DAC5 in water at 18°C, and panels (F–J) display the corresponding structures in 1×PBS at 18°C. Predictions were generated using the DINAMelt Server – Quikfold web tool, and only the structure with the lowest ΔG (Gibbs free energy) was presented.

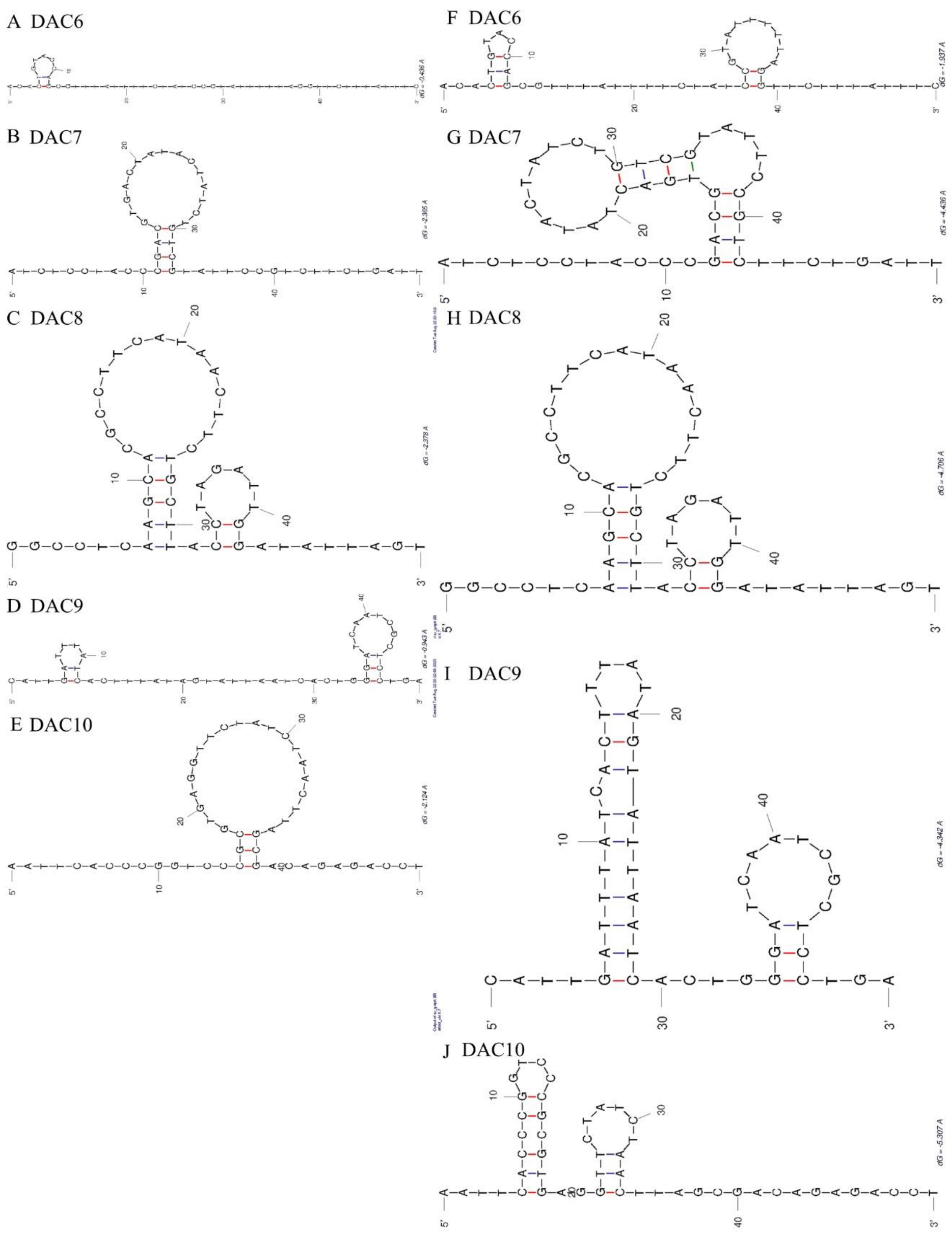


**Fig. S5. Predicted secondary structures of DAC6–DAC10.** Panels (A–E) show the predicted secondary structures of DAC6–DAC10 in water at 18°C, and panels (F–J) display the corresponding structures in 1×PBS at 18°C. Predictions were generated using the DINAMelt Server – Quikfold web tool, and only the structure with the lowest ΔG (Gibbs free energy) was presented.

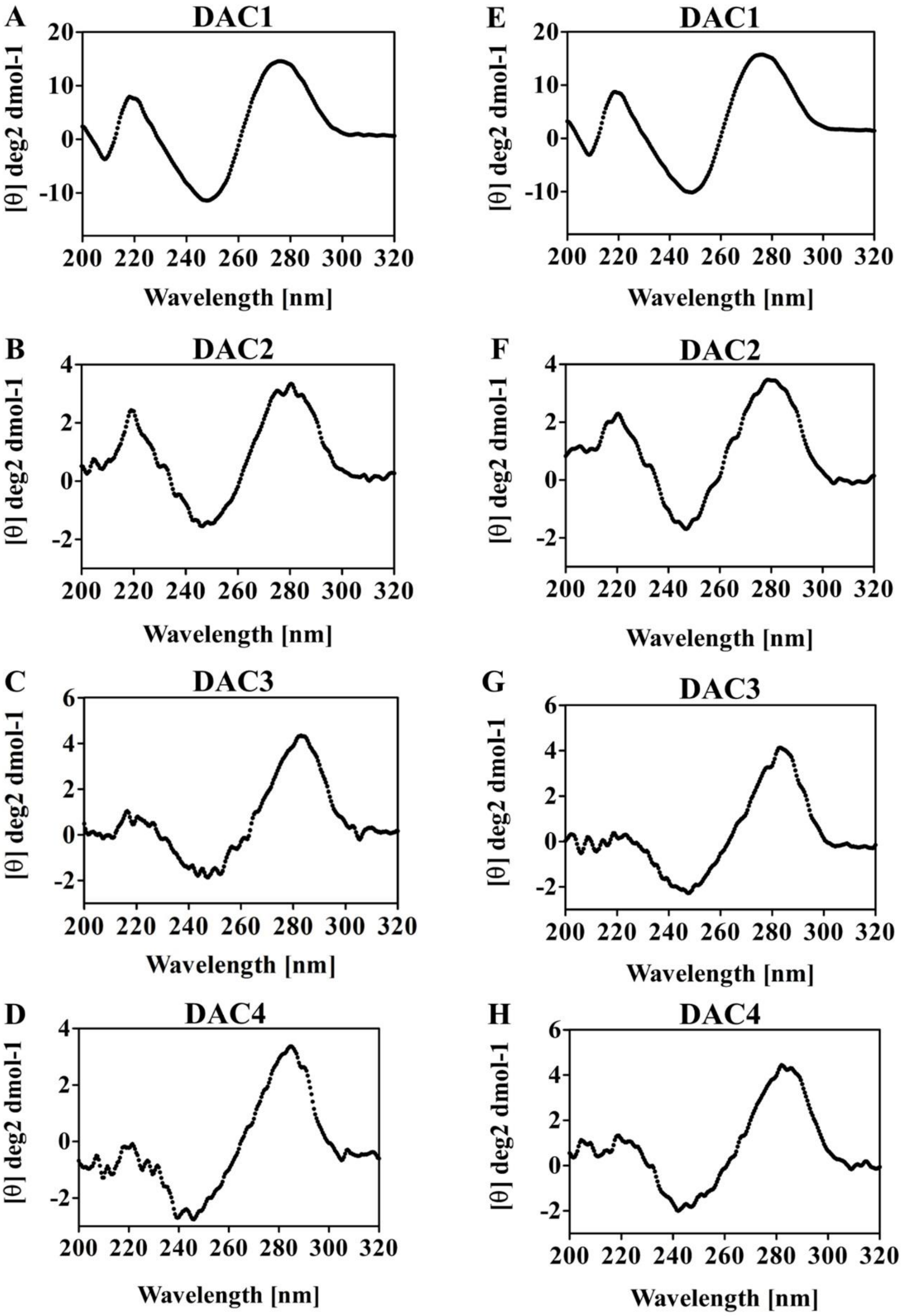


**Fig. S6**. **Secondary structure of DAC1–DAC4.** Panels (A–D) show the CD spectra of DAC1–DAC4 in water at 18°C, while panels (E–H) present the corresponding spectra in 1×PBS at 18°C. All spectra were recorded over the wavelength range of 320–200 nm

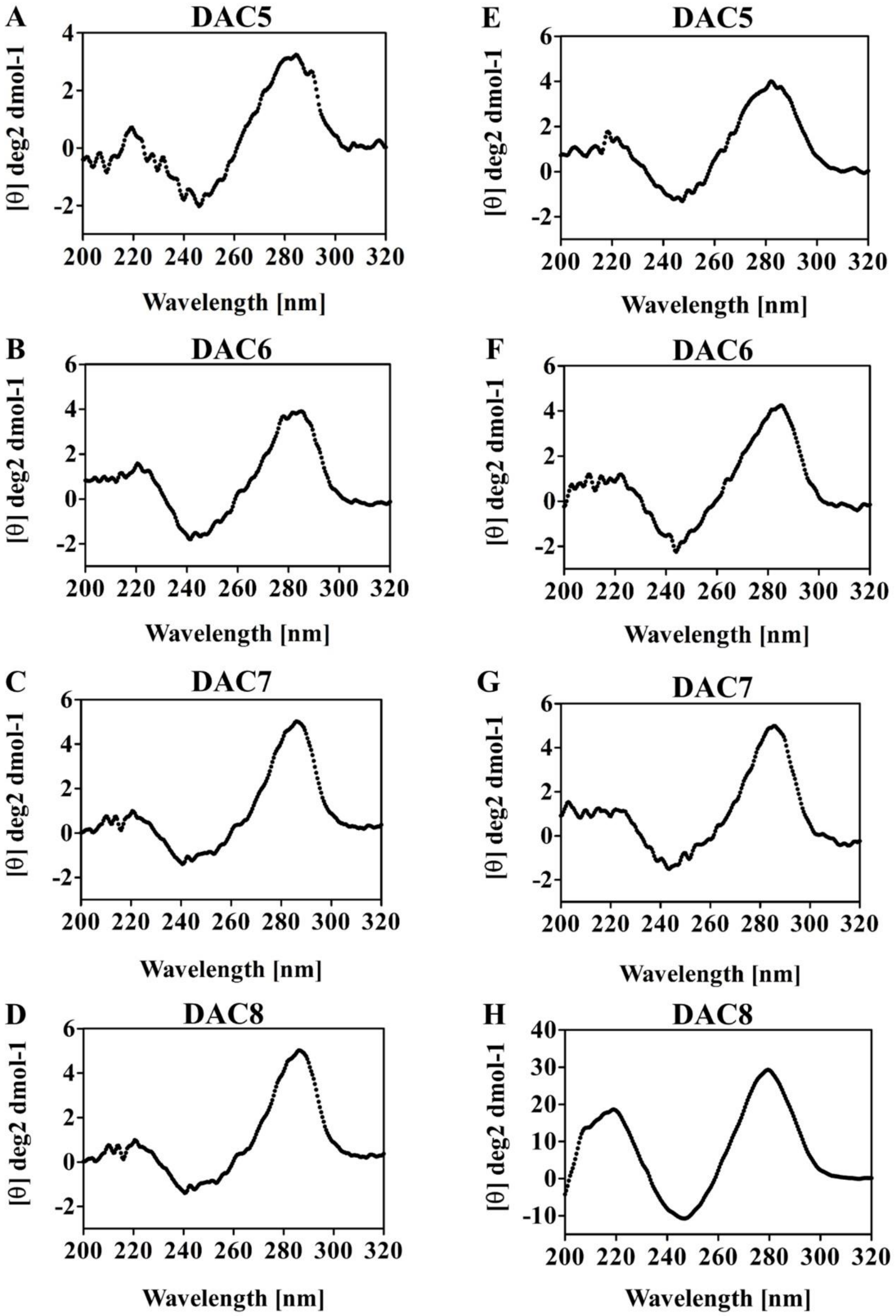


**Fig. S7**. **Secondary structure of DAC5–DAC8.** Panels (A–D) show the CD spectra of DAC5–DAC8 in water at 18°C, while panels (E–H) present the corresponding spectra in 1×PBS at 18°C. All spectra were recorded over the wavelength range of 320–200 nm

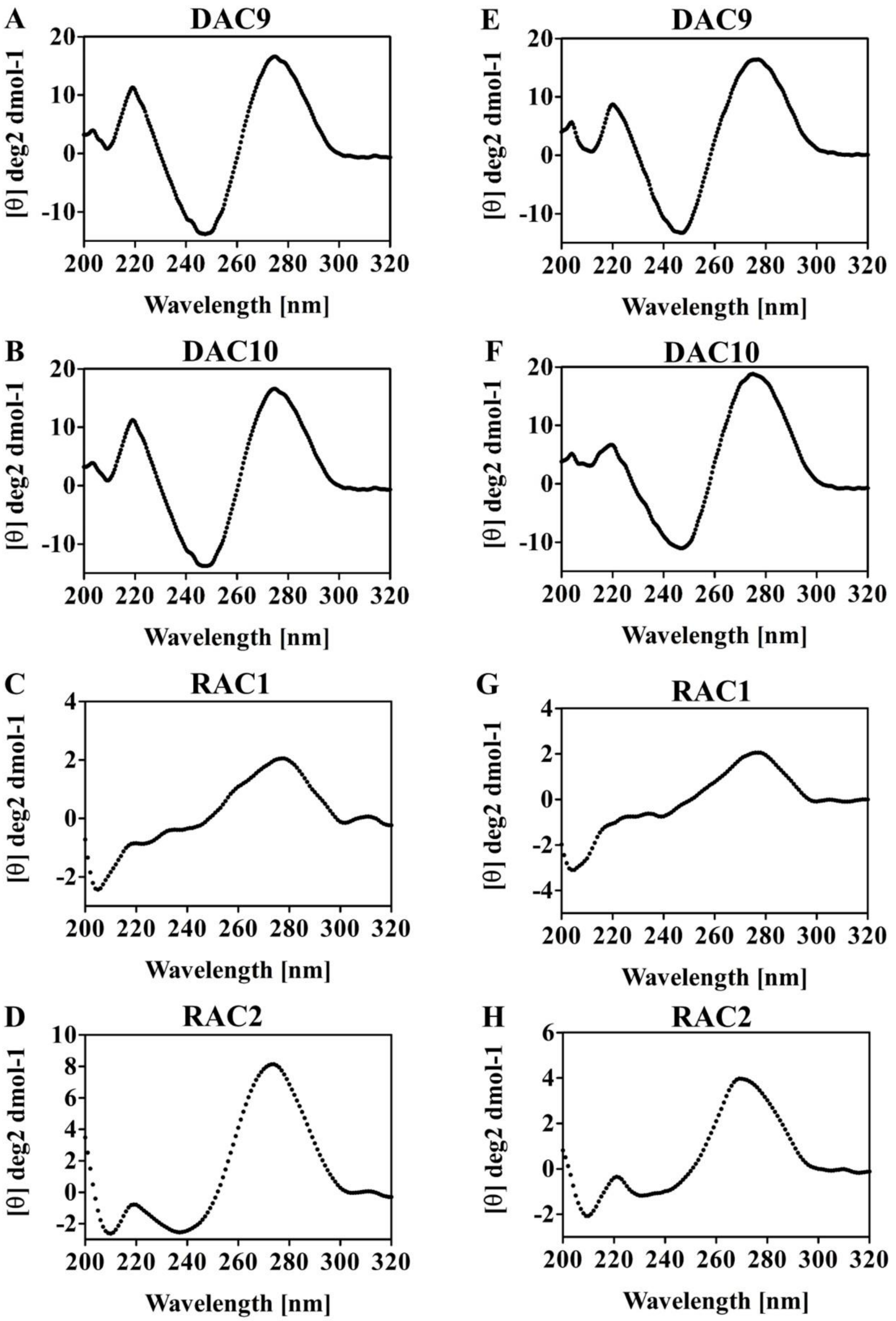


**Fig. S8**. **Secondary structure of DAC9–DAC10 and RAC1-RAC2.** Panels (A–D) show the CD spectra of DAC9–DAC10 and RAC1-RAC2 in water at 18°C, while panels (E–H) present the corresponding spectra in 1×PBS at 18°C. All spectra were recorded over the wavelength range of 320–200 nm

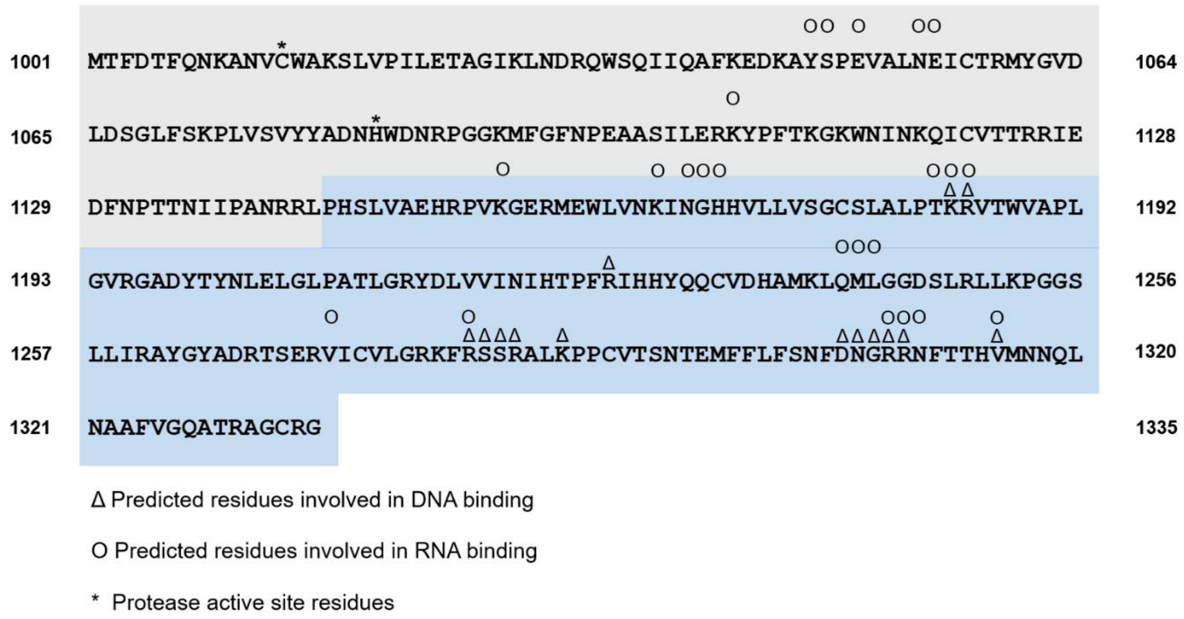


**Fig. S9. Predicted nucleic acid binding regions in the nsP2pro sequence.** The papain-like cysteine protease sequence is displayed with a grey background and the Ftsj methyltransferase (MTase)-like domain is highlighted in blue. The sequence numbering based on the CHIKV polyprotein. Predicted amino acids involved in DNA binding are labeled by a triangle and those predicted to be involved in RNA binding are labeled by a circle. The catalytically active residues of the cysteine protease are labeled by asterisks.

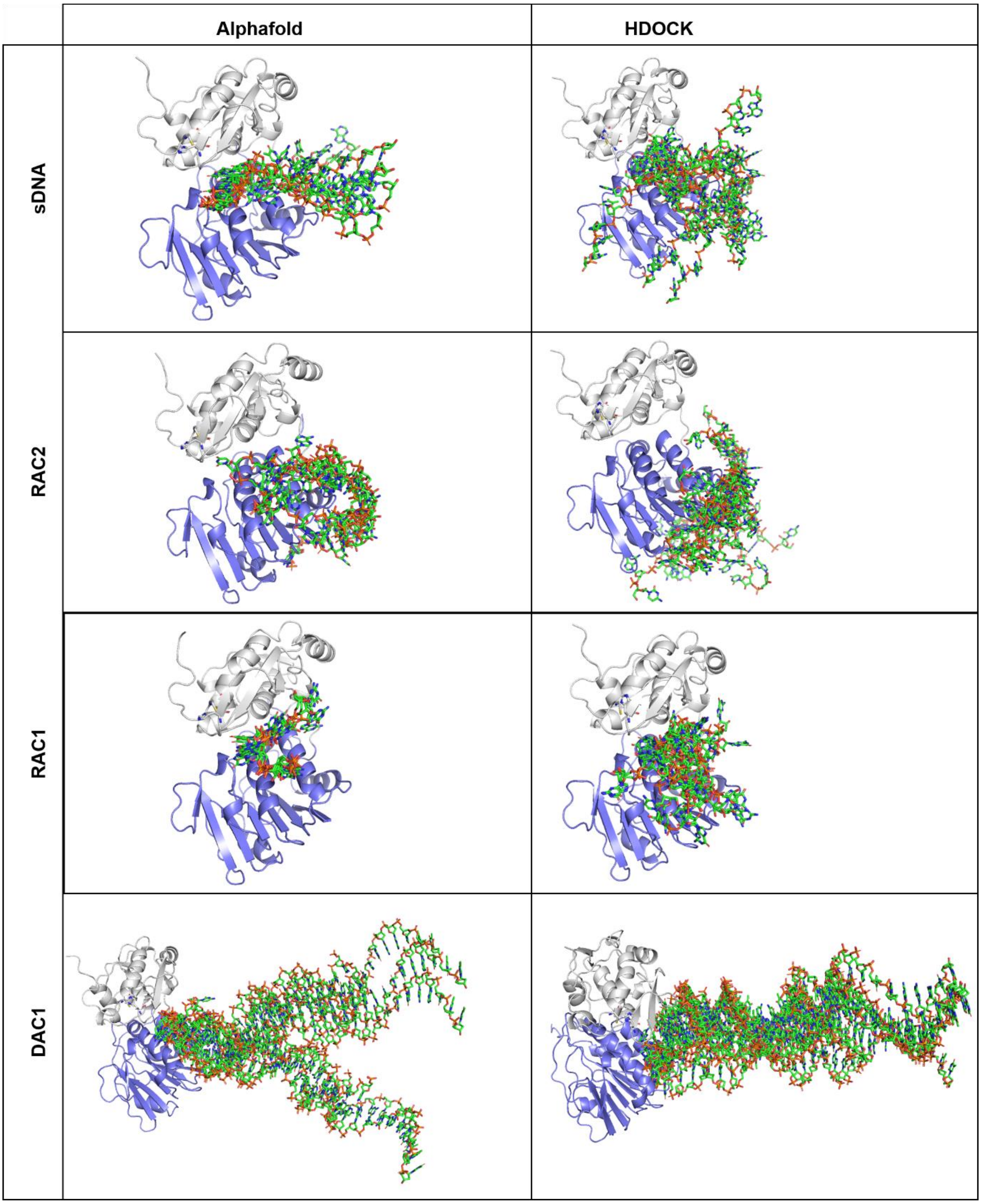


**Fig. S10. Overlay of the predicted nucleic acid positions for AlphaFold models and HDOCK results.** The nsp2$^{pro}$ structures are shown in ribbon view, the papain-like cysteine protease is colored in grey and the Ftsj methyltransferase (MTase)-like domain in blue. The nucleic acids are shown as sticks and in green color. AlphaFold, generated five structures per complex and the overlay of these five structures are shown. The docking with HDOCK generated clusters and the best results of each cluster (depending on the HDOCK score) was used for the overlay.

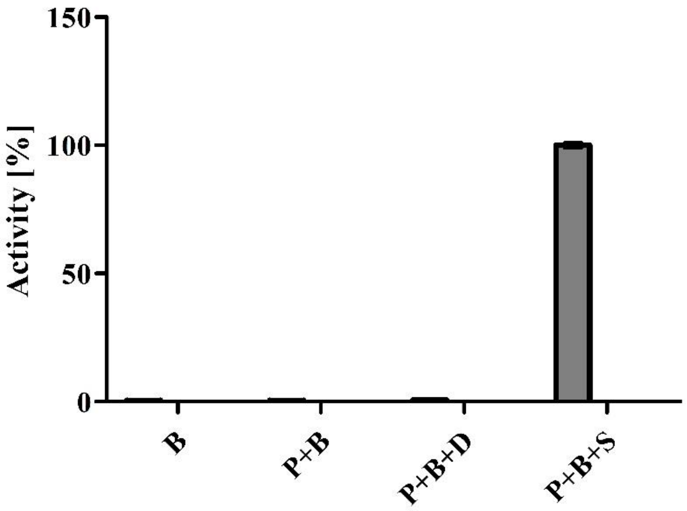


**Fig. S11. Control experiments confirm substrate-specific fluorescence signal in the nsP2$^{pro}$ activity assay**. Fluorescence intensity was measured under four conditions to evaluate potential background signal in the absence of the fluorogenic substrate: (1) buffer alone (B), (2) buffer with CHIKV nsP2$^{pro}$ (P), and (3) buffer with both protease and DNA aptamer DAC8 (D). A positive control containing buffer, protease, and the fluorogenic substrate (Control) was included to represent true enzymatic activity. No significant fluorescence was detected in the absence of the substrate, confirming that background signal from buffer components, protease, or DNA aptamer was negligible.

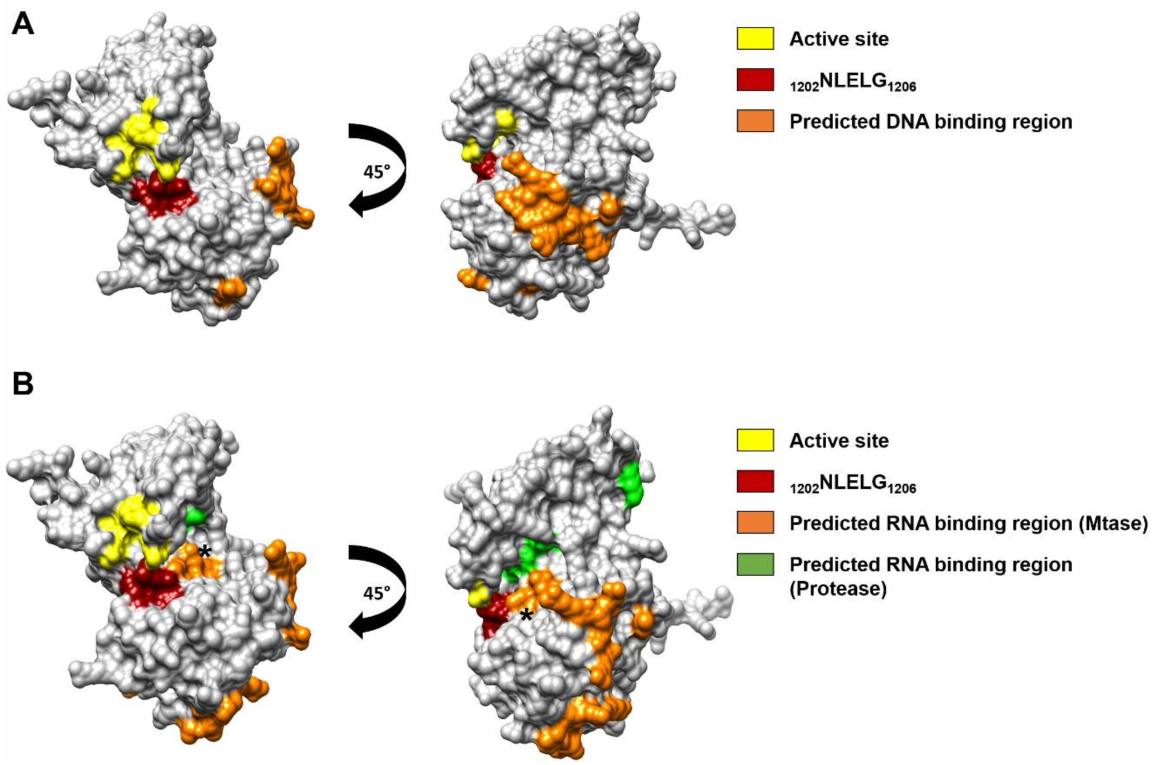


**Fig. S12. Surface view of nsP2$^{pro}$ with labeled predicted nucleic acid binding areas, protease active site and MTase loop $_{1202}$NLELG$_{1206}$.** The nsP2$^{pro}$ active site is shown in close conformation were the protease active site residues Cys1013 and His1083 and the Mtase loop $_{1202}$NLELG$_{1206}$ are nearby. A: Predicted DNA binding region. B: Predicted RNA binding region. Asterisk label the position of $_{1241}$QML$_{1243}$.

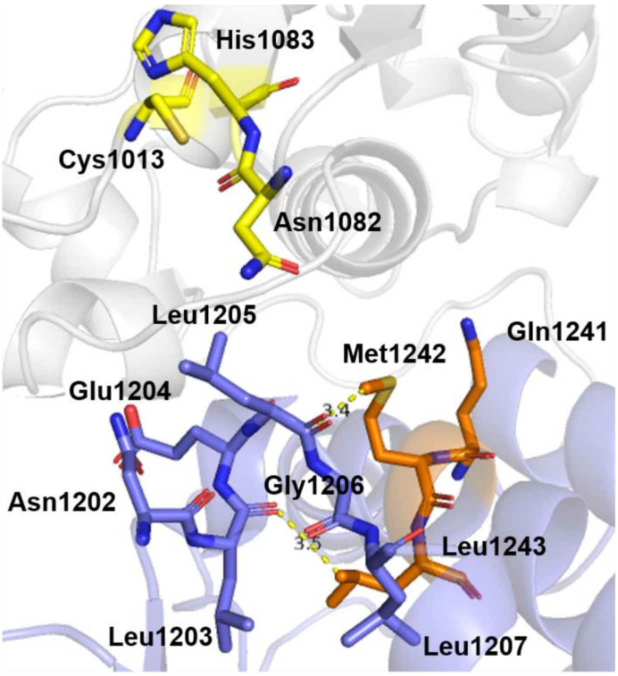


**Fig. S13. Closed conformation of the nsP2$^{pro}$ active site with $_{1202}$NLELG$_{1206}$ and possible $_{1241}$QML$_{1243}$ interactions.** The nsP2$^{pro}$ active site residues are colored in yellow. The Mtase loop $_{1202}$NLELG$_{1206}$ is colored in blue and the Mtase region $_{1241}$QML$_{1243}$ is colored in orange.

**Table S1.** Sequences and properties of the RNAs used in this study. The RNA sequences were derived from the Chikungunya virus (CHIKV) genome (associated GenBank accession: KM673291).

| **RNA** | **Sequence (5'-3')** | **Origin** | **Tm [°C]** | **Manufacturer** |
|---|---|---|---|---|
| **RAC1** | CGTUCG | CHIKV genome | 0.0 | Integrated DNA Technologies (IDT) |
| **RAC2** | CGUCGCUAUA | CHIKV genome | 21.1 | Integrated DNA Technologies (IDT) |
